# A minimal common core and dynamic protein associations orchestrate nuclear mRNP maturation

**DOI:** 10.64898/2026.11.16.767097

**Authors:** PC Fridy, I Abramchuk, AV Ratushny, KE Wei, EY Jacobs, RS Rogers, RA Saleem, LC Aguilar, P Raymond, RL Moritz, D Zenklusen, BT Chait, M Lavallée-Adam, JD Aitchison, MP Rout, M Oeffinger

**Affiliations:** Laboratory of Cellular and Structural Biology, The Rockefeller University, New York, NY, U.S.A; Department of Biochemistry, Microbiology and Immunology, and Ottawa Institute of Systems Biology, University of Ottawa, Ottawa, ON, Canada; Center for Global Infectious Disease Research, Seattle Children’s Research Institute, Seattle, WA, U.S.A; Institute for Systems Biology, Seattle, WA, U.S.A; Laboratory of RNP Biochemistry, Institut de recherches cliniques de Montréal (IRCM), Center for Genetic and Neurological Diseases, Montréal, QC, Canada; Laboratory of Mass Spectrometry and Gaseous Ion Chemistry, The Rockefeller University, New York, NY, U.S.A; Département de biochimie et médecine moléculaire, Université de Montréal, Montréal, QC, Canada; Division of Clinical and Translational Research, McGill University, Montréal, QC, Canada; Department of Pediatrics and Department of Biochemistry (Adjunct), School of Medicine, University of Washington, Seattle, WA, U.S.A

**Keywords:** mRNP assembly, nuclear mRNA maturation, co-transcriptional assembly, mRNA export, nuclear pore basket, chromatin domains, affinity purification–mass spectrometry, AP-RNA-seq, single-mRNP purification, *Saccharomyces cerevisiae*

## Abstract

How nuclear mRNAs are selectively packaged into ribonucleoprotein particles (mRNPs) is fundamental to elucidating the mechanisms that govern gene expression. Here, we interrogate nuclear mRNP maturation in *S. cerevisiae* by combining a proteomic (34 baits) and transcriptomic (18 baits) affinity-purification survey with sequential purification of individual MS2-tagged nuclear transcripts at successive stages of maturation. Contrary to the prevailing models in which nuclear mRNAs are extensively coated with a stable complement of proteins, we find that they share a minimal core of Yra1p and Pab1p, together with a more dynamically associated cap-binding complex. Most other factors associate transiently and transcript-selectively, assembling as pre-assembled, stage-specific modules. The minimal core is invariant to promoter, coding sequence and transcript length, whereas intron class, rather than intron presence alone, determines spliceosome composition. The breadth of factor association further tracks promoter class and proximity to chromatin-domain boundaries, linking mRNP biogenesis to genome organization.

## INTRODUCTION

In eukaryotes, a messenger RNA (mRNA) matures in the nucleus as a dynamic ribonucleoprotein particle (mRNP). As nascent transcripts are synthesized by RNA polymerase II, they are bound co-transcriptionally and post-transcriptionally by successive RNA-binding proteins (RBPs) that carry out 5′ capping, splicing, 3′ cleavage and polyadenylation, packaging, and export through the nuclear pore complex (NPC) for their translation in the cytoplasm^1–4^. mRNP composition shapes every step of an mRNA’s life, from processing efficiency and stability to export competence and cytoplasmic use, and defects in assembly trigger surveillance pathways that degrade aberrant transcripts^1^. Other RNA classes—tRNAs^5^, rRNAs^6,7^ and the sn/snoRNAs^8^—are processed by dedicated machineries acting on transcripts of largely invariant sequence and structure. mRNAs, by contrast, are highly diverse in sequence and structure. The machineries that process them—the cap-binding complex (CBC), the THO–TREX complex that couples elongation to packaging and export, the cleavage and polyadenylation apparatus, SR-like proteins, and the conserved export receptor Mex67p/NXF1—are well characterized in yeast and human cells^1,9,10^, but how they are organized into a single, dynamically remodeled particle as a transcript matures has remained unclear^11^. Central to this is a long-standing question: are all mRNAs matured by a common, ordered set of factors along a single, shared pathway, or is the protein composition of an mRNP tailored to subsets or even individual transcripts?

Studies from many groups have addressed this question, with partly conflicting results. On the one hand, they suggest that mRNP composition is transcript-selective rather than uniform: affinity isolation of individual mRNP proteins coupled to transcriptomics showed that nuclear RBPs bind distinct but overlapping subsets of mRNAs. The shuttling hnRNP-like proteins Nab2p, Npl3p and Hrp1p, for example, each have distinct transcript preferences^12–14^, with Npl3p preferentially binding the intron-containing ribosomal protein (RP) mRNAs^15^. Similarly, the export receptor Mex67p engages mRNAs broadly but is recruited by different RBPs—Nab2p, Npl3p and Yra1p—whose transcript preferences differ, with Nab2p binding shorter and Yra1p longer mRNAs in crosslinking maps^13,16^.

On the other hand, transcriptome-wide crosslinking suggested that Nab2p, Npl3p and Yra1p bind along much of the length of most transcripts^17^, although, given the limited cellular abundance of these proteins, such binding is unlikely to reflect uniform coating of every mRNA molecule. Other observations again point to selectivity: the NPC’s nuclear basket preferentially associates with a subset of RNPs that contain long, intron-poor mRNAs during export^2^, and a survey of RNP composition resolved distinct classes of mRNAs and lncRNAs defined by their bound proteins^16^; some transcripts additionally require the alternative nuclear export factor Crm1p, whose role in mRNA export is best established in metazoa^18–20^. Whether transcript preferences extend beyond these few RBPs to the rest of the maturation machinery remains open. Adding to the complexity, RNA-interactome capture studies revealed an mRNA-bound proteome far larger and more diverse than these classical factors, including many proteins that lack canonical RNA-binding domains and whose function in, and mode of association with, maturing mRNPs remain unknown^21–23^.

A second, related question is how densely an mRNA is coated with proteins. Nuclear mRNAs have been proposed to be extensively coated by RBPs that shield them from nucleases and mark them for export or degradation^17^; alternatively, only a few proteins may be stably bound, with surveillance recognizing aberrant transcripts by other features^24,25^. The question of coating is also tied to packaging: as a long, flexible polymer, an mRNA must be kept from collapsing on itself and organized so that it can be processed, folded and packaged correctly. The mRNA sequence itself contributes to this compaction^26,27^, which in mRNPs is driven by a combination of mRNA–mRNA, protein–mRNA and protein–protein interactions^11,28^. Structural and single-molecule studies have begun to resolve these particles, a few factors at a time. In *S. cerevisiae*, single-molecule imaging quantified the per-mRNP occupancy of export factors on individual transcripts and revealed pronounced molecule-to-molecule heterogeneity^25^. Cryo-electron tomography has revealed compact nuclear mRNP particles, with Yra1p and Yhs7p implicated in their packaging, potentially through the RNA-annealing activity of their positively charged intrinsically disordered regions^29^. In human cells, cryo-electron microscopy (cryo-EM) of the THO–UAP56 complex showed how ATP binding and hydrolysis by UAP56/DDX39B drive deposition of ALYREF onto TREX^30^, and the exon junction complex (EJC) was found to serve as a periodic ALYREF/TREX docking array that compacts the mRNP^28^. What is still missing, however, is a comprehensive, stage-resolved view of the full nuclear mRNP proteome as a transcript matures, and of how that composition relates to the genomic context in which transcription occurs.

These gaps frame four questions. First, is there a universal core—a small set of proteins that remains on transcripts from transcription to export—distinct from factors that associate only with particular transcript classes or processing intermediates? Second, how does composition change as a transcript is capped, spliced, 3′-end processed and exported—by dynamic exchange of individual proteins, or by concerted exchange of defined, pre-assembled complexes? Third, how far do intrinsic transcript features—intron identity and position, promoter class, transcript length—shape the set of RBPs assembled on a given mRNA? And fourth, are these assembly dynamics reflected in the spatial organization of genes in the nucleus, with co-regulated loci associating with specialized structures such as chromosomal interaction domain (CID) boundaries or distinct NPCs?

We address these questions in *S. cerevisiae* using two complementary approaches. First, in a genome-scale survey, we combined affinity purification–mass spectrometry (AP-MS) of 34 mRNP-associated proteins chosen to span the mRNA maturation pathway with RNA sequencing of the same isolations for 18 of them (AP-RNA-seq), mapping protein and transcript co-association across the pathway and identifying candidate core and stage-specific factors. Second, we developed a sequential single-transcript purification that extends single-step MS2-based mRNP isolation: it captures a defined MS2-tagged mRNA at successive nuclear maturation stages and allows measurement of the composition and copy number of its mRNP^31–33^. Earlier binding surveys could identify broadly associated factors but could not determine how many copies of a factor occupy a single mRNP, resolve a particle at a defined stage, or distinguish factors that join individually from those that join as complexes; the single-transcript approach addresses these limitations. Together, the two approaches show that only a few proteins—Yra1p and Pab1p, together with the more dynamically associated CBC—persist across all maturation stages, whereas most other factors are organized into discrete, stage-specific complexes that load onto the transcript as pre-assembled modules rather than as individual proteins. In the survey, the breadth of factor association further tracks transcript class, promoter class and the spatial organization of the genome. These data provide an integrated genome-scale and single-particle view of nuclear mRNP biogenesis and define the minimal protein core of a maturing mRNA.

## RESULTS

### Survey of the nuclear mRNP network reveals a modular, stage-defined architecture

To investigate the mRNP assembly landscape, we selected 34 proteins with established roles in nuclear mRNA maturation, spanning eight functional groups spread along the pathway: cap binding (CBC), co-transcriptional assembly/THO/TREX, splicing, 3′-end processing, RNA surveillance, TREX-2/NPC, nuclear export, and cytoplasmic factors (Figure 1A). Besides representing distinct stages, the selected proteins included factors essential for general mRNA biogenesis, functional paralogs that probe redundancy, and factors with documented transcript-specific roles such as splicing regulators and localization factors, as well as a nuclear mRNP-associated protein of poorly defined function (Yhs7p).

**Figure 1.**
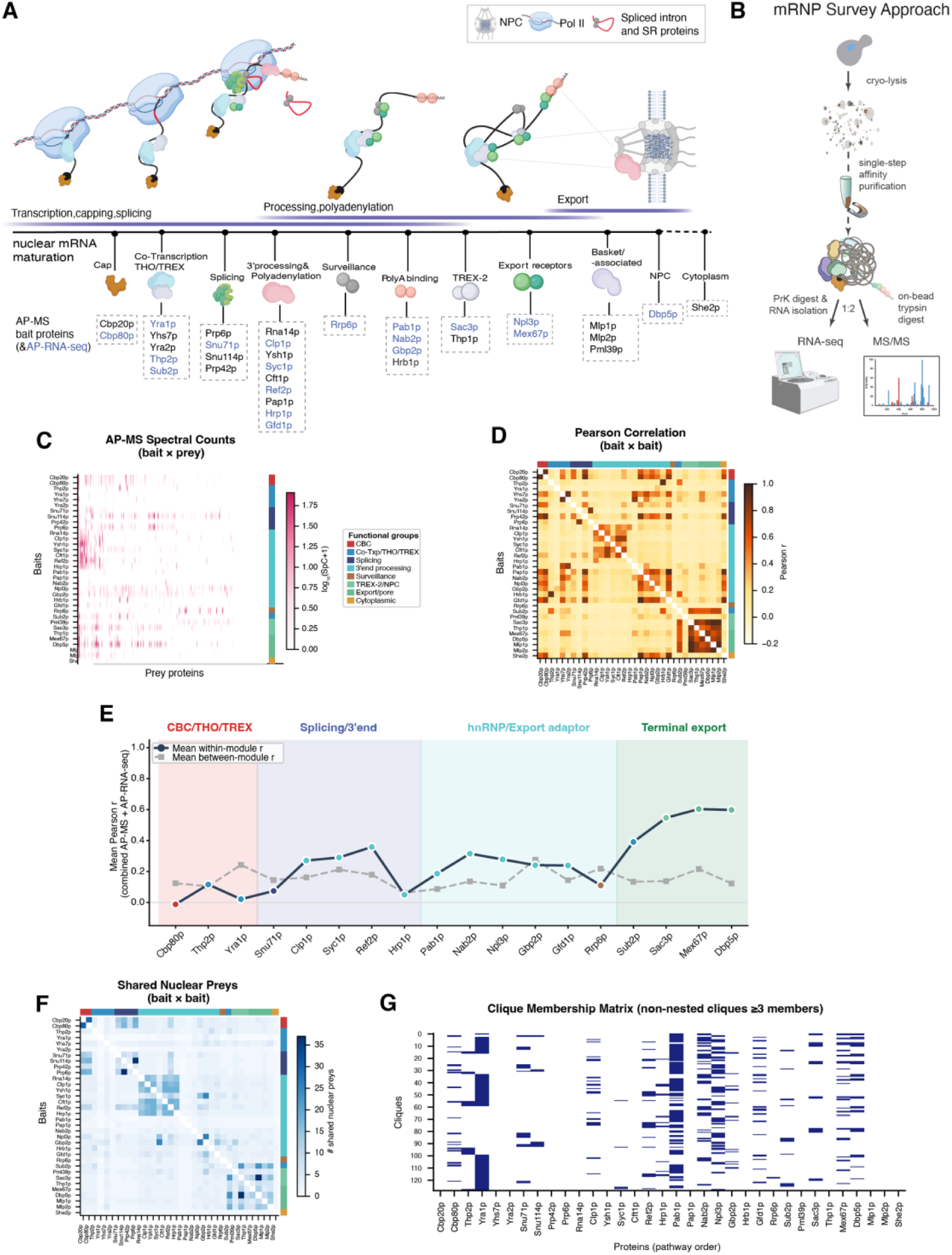
Survey of the nuclear mRNP network reveals a modular, stage-defined architecture. **(A)** Experimental design schematic. The 34 nuclear mRNA biogenesis factors spanning co-transcriptional assembly (CBC, THO/TREX), splicing, 3′-end processing, RNA surveillance, TREX-2/NPC, and export used as baits in affinity purification–mass spectrometry (AP-MS). Baits are color-coded by functional stage throughout all figures. **(B)** mRNP survey workflow. Yeast expressing each PrA-tagged bait were cryogenically lysed; bait-associated complexes were affinity-purified on IgG-conjugated magnetic beads and the eluate split for parallel analysis of co-purified protein (AP-MS) and RNA (AP-RNA-seq) from the same preparation. **(C)** AP-MS spectral counts. log₁₀(SpC+1) for nuclear-filtered prey proteins (columns, hierarchical clustering order) across the bait proteins (rows, pathway order). Functional-group color bar at right; white = not detected, dark pink = highest. **(D)** Bait–bait Pearson correlation matrix of AP-MS prey profiles. Pairwise Pearson correlation of spectral-count enrichment profiles across all shared nuclear preys, baits ordered by maturation stage. Major correlation blocks correspond to the four functional modules, CBC/THO–TREX, splicing/3′-end processing, hnRNP/export adaptor and terminal export (TREX-2/Mex67p); the same matrix in clustering order is shown in Supplemental Figure 2B. Color scale: dark amber = high positive correlation. **(E)** Within-versus between-module co-purification correlation per bait, for the 18 baits profiled by both AP-MS and AP-RNA-seq. For each bait, the mean combined (AP-MS + AP-RNA-seq) Pearson correlation with other members of its module (within; filled circles, solid line) versus with baits in other modules (between; gray squares, dashed line). Baits are ordered by module and shaded by module membership (CBC/THO–TREX, Splicing/3′-end, hnRNP/Export adaptor, Terminal export); within-module correlation is significantly higher than between-module correlation (one-sided Mann–Whitney U test over bait pairs, p = 0.016). **(F)** Shared nuclear prey matrix. Number of nuclear-localized prey proteins co-purified in common by each pair of baits (bait × bait, pathway order); color scale indicates shared-prey count. **(G)** Clique membership matrix. Rows: the 129 non-nested cliques (≥3 members) in the undirected bait co-detection graph; columns: bait proteins in pathway order. Filled cells indicate membership.

Each factor was tagged at its endogenous locus with a Protein A (PrA) epitope on its C-terminus as the sole source of the protein^34^. Cells were grown to mid-log phase, snap-frozen, and cryogenically milled, a process demonstrated to efficiently access all the cellular contents while preserving intact mRNP complexes^35^. Bait complexes were captured on IgG-conjugated magnetic beads under buffer conditions optimized for each bait, as confirmed by Coomassie-stained SDS-PAGE (Supplemental Figure 1A; Supplemental Table 3F), and each eluate was split for parallel mass spectrometry (AP-MS) and RNA sequencing (AP-RNA-seq), so that protein and RNA co-purification were analyzed from the same sample (Figure 1B). For AP-MS, we restricted the analysis to nuclear-annotated proteins to focus on the nuclear phase of mRNP maturation; this set aside ∼11% of spectral counts, largely translation factors and cytoplasmic mRNA-binding partners of the shuttling baits (e.g., Cdc33p, Sup35p, Ded1p).

Co-purification enrichment across all 34 AP-MS baits showed distinct profiles across functional bait groups that recapitulated their known functional organization (Figure 1C; Supplemental Figure 2A, E, F). Factors acting at the same step co-purified together: the CBC subunits Cbp20p and Cbp80p; the 3′-end cleavage and polyadenylation factors Rna14p, Clp1p, Ysh1p, Hrp1p, Cft1p and Ref2p; and the co-transcriptional THO/TREX components Thp2p, Sub2p, and Yra1p. In addition, the Yhs7p bait co-purified Yra1p. Notably, the paralogous pairs Gbp2p/Hrb1p and Ysh1p/Syc1p did not reciprocally co-isolate one another, consistent with their membership in distinct complexes—Ysh1p in the CPF complex that processes coding-mRNA 3′ ends, and Syc1p in the APT complex that forms sn/snoRNA 3′ ends^36–38^. The separation of these paralogs into distinct interactomes demonstrated the specificity of our approach (Figure 1C, D, F).

Bait–bait Pearson correlation of the co-purification profiles, with baits ordered by maturation stage, revealed four broad modules of bait proteins with similar profiles that follow the pathway’s direction: early biogenesis (CBC/Co-Txp/THO/TREX), mid-pathway processing (splicing/3′-end), late nuclear (hnRNP/export adaptor), and terminal export (TREX-2/Mex67p) (Figure 1E; Supplemental Figure 2B). These modules remained distinct when protein and RNA co-purification were combined. Across the 18 baits profiled by both AP-MS and AP-RNA-seq (Figure 1B), within-module correlation was significantly higher than between-module correlation (one-sided Mann–Whitney U test over bait pairs, p = 0.016), with the CBC/THO–TREX as the only exception (Figure 1E). This exception likely reflects that two of its three baits, Cbp80p and Yra1p, are not restricted to the earliest stage but span the full maturation network, therefore correlating more strongly with later modules than with each other.

Pairwise shared-prey counts—i.e., the number of nuclear preys co-purified by each bait pair, across all 34 baits—were highest within functional groups and decreased the further apart two baits functioned along the pathway, reinforcing the modular organization seen by correlation (Figure 1F). Distances across the bait–prey network independently confirmed this organization: when each bait–prey edge was weighted by co-purification strength (edge cost inversely proportional to spectral-count enrichment), baits of the same stage were separated by shorter distances than baits of different stages (Supplemental Figure 3A–D). Beyond this modularity, the network also reflected the order of the pathway. In a directed network (i.e., edges A→B mark detection of B as a prey when A is the bait), connectivity was asymmetric and followed the pathway’s direction: early baits (THO/TREX) were recovered as prey by downstream baits, while terminal export factors were rarely detected by upstream baits (Supplemental Figure 3E). Betweenness centrality, which measures how often a protein lies on the shortest paths between other proteins in the network, identified the THO/TREX components Yra1p, Thp2p and Sub2p as the main links between functional groups (Supplemental Figure 3F), consistent with TREX coupling transcription to export.

To resolve the modular architecture further, we applied clique analysis to identify fully connected sets of baits—groups that mutually co-purify one another and so likely share complexes. This identified 129 distinct non-nested cliques across the 34 baits, consistent with a rich subcomplex structure within each stage (Figure 1G). Baits differed widely in how many distinct nuclear preys they recovered, from 1 to 61 (median 21; Supplemental Figure 1B, C).

We next asked the converse: which proteins are recovered as preys by the greatest number of baits—i.e., the most broadly connected nodes of the co-purification network (Supplemental Figure 3G). The poly(A)-binding protein Pab1p was detected in essentially every purification (31 of 34). Similarly, Yra1p was present in the large majority (26) and Npl3p in half (17), mainly with CBC, splicing and 3′-end processing baits^14,17,39,40^, both far more broadly than any other factor (at most 8; Supplemental Figure 2C, D). These three participated in the most cliques, several of which also contained the CBC. Moreover, the CBC (Cbp80p, Cbp20p) and Cdc33p (eIF4E), which is also partly nuclear^41^, were recovered by mutually exclusive sets of baits (the CBC by its own subunits and Npl3p; Cdc33p by Pab1p, Nab2p, Gbp2p, Gfd1p and Dbp5p), suggesting that the exchange of cap-binding proteins, generally thought to occur during or after export^42^, may already begin at late, export-associated nuclear stages. Proteins recovered in only a few purifications were predominantly functional paralogs that occupy distinct complexes, such as Syc1p and Gbp2p (above) and Yra2p, or factors acting specifically in late export (e.g., Mex67p, Mlp1p, Sac3p), likely reflecting specialization rather than redundancy (Supplemental Figure 1D). The broadly connected factors were not confined to any one stage but spanned the full biogenesis network, suggesting they are shared components of maturing mRNPs across stages rather than stage-specific factors.

The co-purification subnetwork of Pml39p, a nuclear basket-associated protein^43^, illustrates such stage-specific topology: Pml39p associated with TREX-2, the nuclear basket and export factors and is set apart from the earlier processing modules, placing it functionally at the interface where maturing mRNPs are passed to the export machinery (Supplemental Figure 3H). Notably, because TREX-2 and the nuclear basket occupy distinct sites at the NPC^44^, this co-purification most likely reflects the shared engagement of Pml39p and TREX-2 with the exporting mRNP as it moves from the basket into the central channel of the NPC. Consistent with this, the TREX-2 bait Sac3p itself co-purified essentially the entire NPC, the nuclear-basket proteins Mlp1p and Mlp2p, and its TREX-2 partners Sus1p, Cdc31p and Sem1p, placing the complex physically at the NPC^44^ (Soni *et al*., manuscript in preparation).

Taken together, the AP-MS survey revealed two organizing principles of the network. First, the machinery resolves into discrete complexes that act in delineated steps: from cap binding through THO/TREX, splicing and 3′-end processing to export, the baits of each complex shared more similar co-purification profiles with one another than with those of adjacent steps (Figure 1C, D, F, G). Maturing mRNPs are thus built from separable, stage-specific modules rather than one continuous assembly. Second, a small set of factors, Pab1p, Yra1p and Npl3p, together with the CBC, pervade the network, spanning most maturation stages, and represent candidate members of a common core. Individual stage mRNP complexes and adaptor preferences have been described before; our survey shows how these modules and the broadly shared factors are organized relative to one another across mRNA maturation. We next asked whether this organization is reflected in the RNAs these factors co-purify.

### AP-RNA-seq identifies transcript classes with distinct co-purification profiles

To ask with which transcripts these factors associate, we performed AP-RNA-seq on 18 of the 34 baits, chosen to represent the stages of the pathway (Figure 2A). Co-purification enrichment scores were computed for 2,452 expressed coding genes (Supplemental Figure 4A) and hierarchically clustered, resolving eight transcript clusters (k = 8; Supplemental Figure 4B). Each cluster therefore groups transcripts co-purified by a similar set of factors; we named the clusters after the baits that enriched them most strongly: C1 CBC/splicing/Npl3p, C2 hnRNP/poly(A), C3 THO/TREX, C4 Hrp1p, C5 3′-end processing, C6 TREX-2, C7 Gfd1p/CBC and C8 export (Figure 2B). Co-purification was highly non-uniform: distinct groups of mRNAs associated preferentially with different subsets of factors (Figure 2A; Supplemental Figure 5A), suggesting that across the transcript population, mRNP composition is not uniform but differs between transcript classes. Moreover, the mean enrichment profiles of the clusters shifted progressively from early-stage (CBC, THO/TREX) to late-stage (export) baits (Figure 2B). Different transcript classes are thus preferentially captured at different stages of maturation, possibly reflecting the step at which each class resides longest.

**Figure 2.**
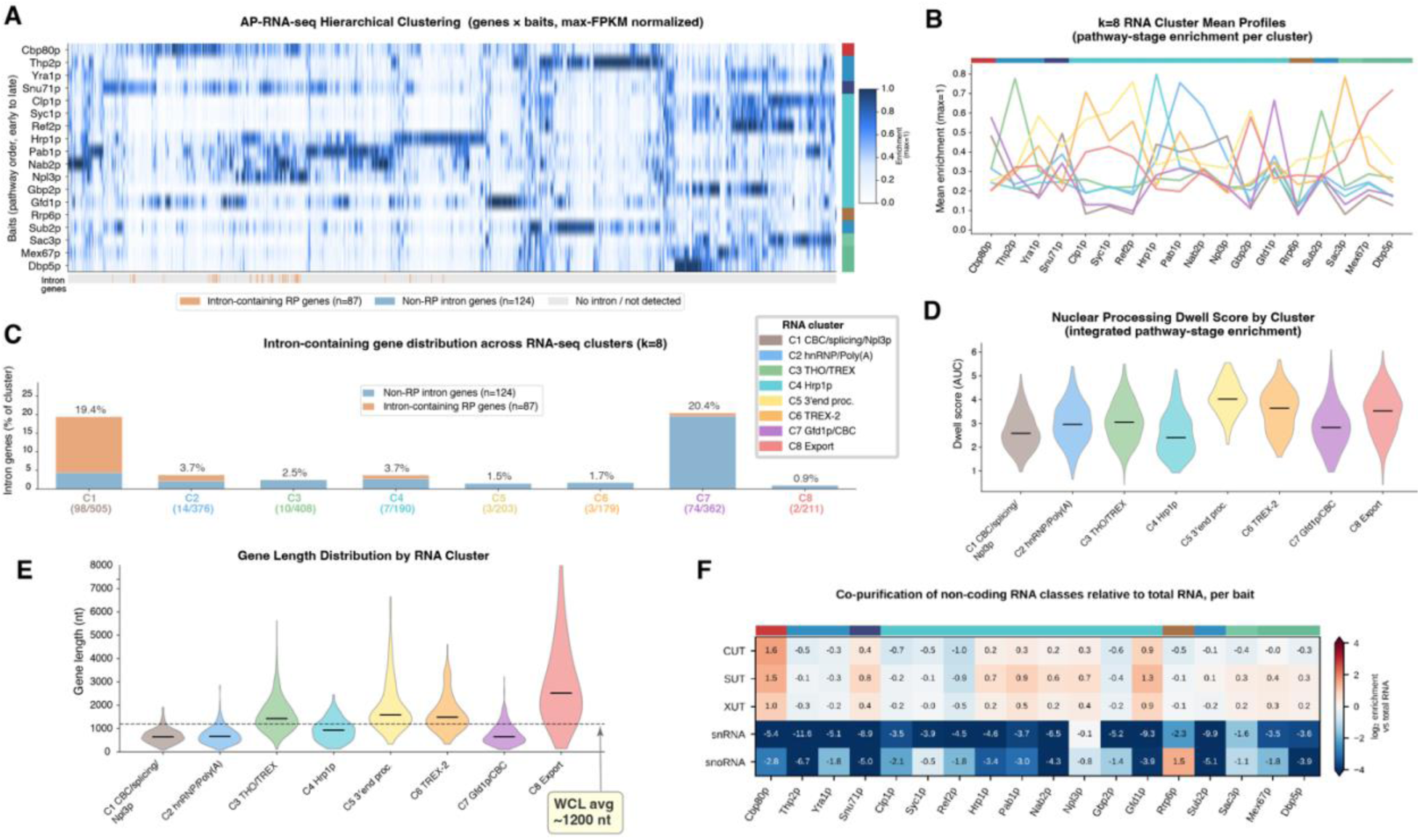
Transcript co-enrichment landscape across nuclear mRNA biogenesis stages. **(A)** Hierarchical clustering of AP-RNA-seq co-enriched transcripts. Max-FPKM-normalized enrichment across 18 baits (rows, pathway order: CBC at top, nuclear export at bottom) for 2,434 co-enriched coding ORFs (columns). Color: dark blue = enriched; white = absent. A three-color strip below the heatmap marks intron-containing genes: intron-containing ribosomal protein (RP) genes (terracotta), non-RP intron-containing genes (dusty blue), non-intron genes (light gray). **(B)** k = 8 cluster mean enrichment profiles. Mean max-normalized enrichment per bait (pathway order) for each transcript cluster (C1–C8). **(C)** Intron-containing gene distribution across k = 8 transcript clusters. Stacked bars showing the percentage of each cluster’s membership that contains spliceosomal introns, split into non-RP intron-containing genes (dusty blue) and RP genes (terracotta). Numbers in parentheses: intron-gene count / total cluster size. χ² = 436.7, p = 2.3×10⁻⁸⁴. **(D)** Nuclear processing dwell score by transcript cluster. Distribution of dwell scores (sum, over the eight pathway stages, of each gene’s highest mean enrichment; see Methods) per k = 8 cluster; higher dwell reflects broader biogenesis-stage co-association. Horizontal line: median. **(E)** Gene length distributions by transcript cluster. ORF lengths (nt) per k = 8 cluster. Horizontal line: median; dashed reference: approximate whole-transcriptome mean (∼1,200 nt). Transcripts of the export (C8), 3′-end processing (C5), TREX-2 (C6) and THO/TREX (C3) clusters are longer than those of the other clusters (one-sided Mann–Whitney U vs. all other clusters, p < 10⁻¹⁸). **(F)** Enrichment of non-coding RNA classes with each bait relative to total RNA. For each bait, the ratio of the median FPKM of CUTs, SUTs, XUTs, snRNAs and snoRNAs to that of coding mRNAs, relative to the same ratio in total RNA from wild-type cells (log₂; 0 = proportion as in total RNA; color scale capped at ±4). Baits in pathway order.

We next asked whether transcripts grouped by co-purification also share biological functions. Gene Ontology (GO) analysis per bait (Supplemental Figure 4C) and per cluster (Supplemental Figure 4D) showed that five clusters were enriched for functionally related transcripts: cytoplasmic translation and ribosome biogenesis in C1, mitochondrial translation in C2, chromatin organization and DNA metabolism in C3, amino-acid metabolism and tRNA aminoacylation in C5, and carbohydrate metabolism in C6; C4, C7 and C8 showed little GO-slim enrichment. Transcripts of related function therefore tended to co-purify with similar sets of maturation factors.

Beyond function, the clusters also differed in gene architecture. Most strikingly, intron-containing transcripts segregated into two clusters, C1 and C7 (Figure 2C). Although fewer than 5% of yeast genes carry introns, intron-containing genes made up 19.4% of C1, which was enriched by the splicing bait Snu71p together with Npl3p and Cbp80p, and 20.4% of C7, which was associated with Gfd1p, a poorly characterized factor that co-purified 3′-end processing and export-interface components^45,46^, and with Cbp80p. In yeast, intron-containing genes fall broadly into two mechanistically distinct groups. Ribosomal protein (RP) genes, which are heavily intron-bearing (66.9% carry introns, vs. 4.6% genome-wide), concentrated almost entirely in C1 (89.2% of RP genes), likely reflecting prolonged co-transcriptional association of the spliceosome with RP pre-mRNAs rather than surveillance-related retention. Non-RP intron-containing genes (n = 124) instead resided mainly in C7, which captured 57% of them (χ² = 436.7, p = 2.3×10⁻⁸⁴; Figure 2B, C). Every other cluster fell below 4% intron content.

Although both intron groups showed short nuclear dwell times (a measure of how long transcripts remain engaged with the processing machinery, defined below; Figure 2D), their differential association suggests distinct splicing dynamics: RP pre-mRNAs of C1 remain engaged with the splicing apparatus, whereas the non-RP intron transcripts of C7 have been released by the splicing machinery and instead associate with Gfd1p^45,46^, indicating that the two classes dwell at different stages of splicing rather than being spliced at different overall rates. Notably, the two intron classes carry equally strong, canonical splice sites—essentially all are GT–AG with comparable 5′-splice-site (GUAUGU) and branchpoint (UACUAAC) consensus matches—but differ chiefly in length, RP introns being ∼4-fold longer (median 407 vs. 100 nt) and more tightly 5′-proximal; their distinct factor associations therefore reflect intron architecture and gene context rather than splice-site strength^47,48^. Intron architecture and gene context thus shape splicing-factor co-purification specificity, and our AP-RNA-seq survey captured biologically meaningful RBP–transcript preferences.

Transcripts also differed in length between clusters: the export cluster C8 contained the longest transcripts, whereas C1, C7 and C2 contained the shortest (Figure 2E), the short C1 transcripts being consistent with the compact size of most ribosomal protein genes. To resolve this at the level of individual baits, we computed Spearman correlations between gene length and enrichment for each bait (Supplemental Figures 4E, 5B). Cap-binding and poly(A)/hnRNP baits (Pab1p, Cbp80p, Hrp1p, Npl3p, Gfd1p and Nab2p), as well as Snu71p, were preferentially enriched on shorter transcripts, whereas 3′-end processing (Clp1p, Ref2p, Syc1p), export (Gbp2p, Sac3p, Mex67p, Dbp5p) and surveillance (Rrp6p) factors, together with Yra1p, were enriched on longer transcripts, as were, to a lesser extent, the THO/TREX components Thp2p and Sub2p, as also suggested by crosslinking maps in which Nab2p preferentially bound short and Yra1p long mRNAs^17^. The preference of Yra1p for longer transcripts agrees with the 5′-to-3′ increase in TREX occupancy along genes, on which the expression of long transcripts specifically depends^49^. Longer transcripts thus become progressively enriched for late-stage and export factors, consistent with their longer residence in the maturation pathway and with the preferential routing of long, intron-poor transcripts through the nuclear basket in yeast^2^ (Supplemental Figure 4F; Supplemental Figure 5C).

The enrichment of longer transcripts for late-stage factors raised the question of how long transcripts remain engaged with the maturation machinery. To estimate this, we derived a nuclear processing dwell score for each gene, based on the integrated co-purification enrichment across all pathway baits ordered by stage (Figure 2D). Dwell scores were highest for the late-stage clusters—the 3′-end processing cluster C5 and the export/TREX-2 clusters, C8 and C6, respectively—consistent with prolonged mRNP maturation before transit through the NPC (Supplemental Figure 4G; per bait, Supplemental Figure 5D).

Although the clusters were built from coding transcripts, the AP-RNA-seq data also captured non-coding RNAs. Relative to their proportions in total RNA, cryptic non-coding transcripts (CUTs, SUTs, XUTs) were enriched most with Cbp80p and Gfd1p (Figure 2F), consistent with cap binding of all Pol II transcripts and the early nuclear decay of CUTs^50^. The nuclear surveillance factor Rrp6p co-purified preferentially with snoRNAs (Figure 2F), consistent with its role in nuclear RNA processing and quality control^50,51^. Npl3p showed weaker but clear snRNA/snoRNA co-purification, whereas Nab2p co-purified 3′-extended snoRNA sequences much more strongly than mature snoRNAs, consistent with its binding to polyadenylated snoRNA precursors that are matured by the exosome and Rrp6p^52,53^. Thus, all factors surveyed predominantly engaged polyadenylated coding transcripts, and only a few were additionally enriched for particular non-coding RNA classes.

Consistent with their broad protein-level connectivity, Pab1p and Yra1p, together with Nab2p, co-purified transcripts across all eight RNA clusters, in contrast to stage-specific factors such as Thp2p, Snu71p and Rrp6p (Supplemental Figure 3I). Nab2p and Pab1p co-purified largely overlapping transcript sets (73% of Nab2p-enriched transcripts were also enriched with Pab1p; 1.3-fold above chance) and showed reproducible transcript preferences, with Pab1p more strongly enriched on TREX-2 cluster (C6) and Nab2p on THO/TREX cluster (C3) transcripts (Supplemental Figure 5E); unlike Pab1p, however, Nab2p was recovered as a prey in only a few purifications (6 of 34, versus 31 for Pab1p). Taken together, the proteomic and transcriptomic readouts of the survey converged on a processing scheme in which a small set of pervasive factors is accompanied by a much larger, stage- and transcript-specific cohort.

To determine whether protein interactions (AP-MS) and transcript co-purification (AP-RNA-seq) provide a consistent picture of pathway organization, we examined the combined correlation matrix of the 18 shared baits (Figure 3A). Hierarchical clustering of this matrix, which does not depend on pathway order, separated the baits into two strongly supported principal branches, co-transcriptional and export-directed (Supplemental Figure 6A). Cutting the tree into four groups reproduced the four modules defined above (Figure 1E), which were also resolved by PCA and multi-omics network construction (Supplemental Figure 6B–D). Within this modular organization, Thp2p and Sub2p resolved as a distinct but loosely joined pair, their exclusive grouping being the least bootstrap-supported of the four groups (AU ≈ 80; Supplemental Figure 6A), consistent with the staged assembly of TREX, in which Thp2p is recruited co-transcriptionally as part of THO and Sub2p is subsequently transferred to the nascent RNA^54^. Their ensemble co-purification profiles accordingly straddled the co-transcriptional and export-directed groups. The two readouts were concordant mainly within functional groups: bait pairs of the same complex, such as Sub2p–Thp2p, Clp1p–Ref2p and Mex67p–Dbp5p, correlated strongly in both, whereas several factors that shared transcripts did not co-purify each other’s proteins (e.g., Yra1p with Mex67p and Dbp5p). The hnRNP/poly(A) group was the least concordant, largely because Gbp2p, which co-purified Nab2p and Npl3p, associated with a distinct, generally longer set of transcripts, and Hrp1p correlated weakly with the other group members in both readouts (Supplemental Figure 7A–D). Protein interaction and transcript co-purification are thus complementary readouts of the same biogenesis network. We next asked whether this organization is mirrored by where the corresponding genes reside within the nucleus.

**Figure 3.**
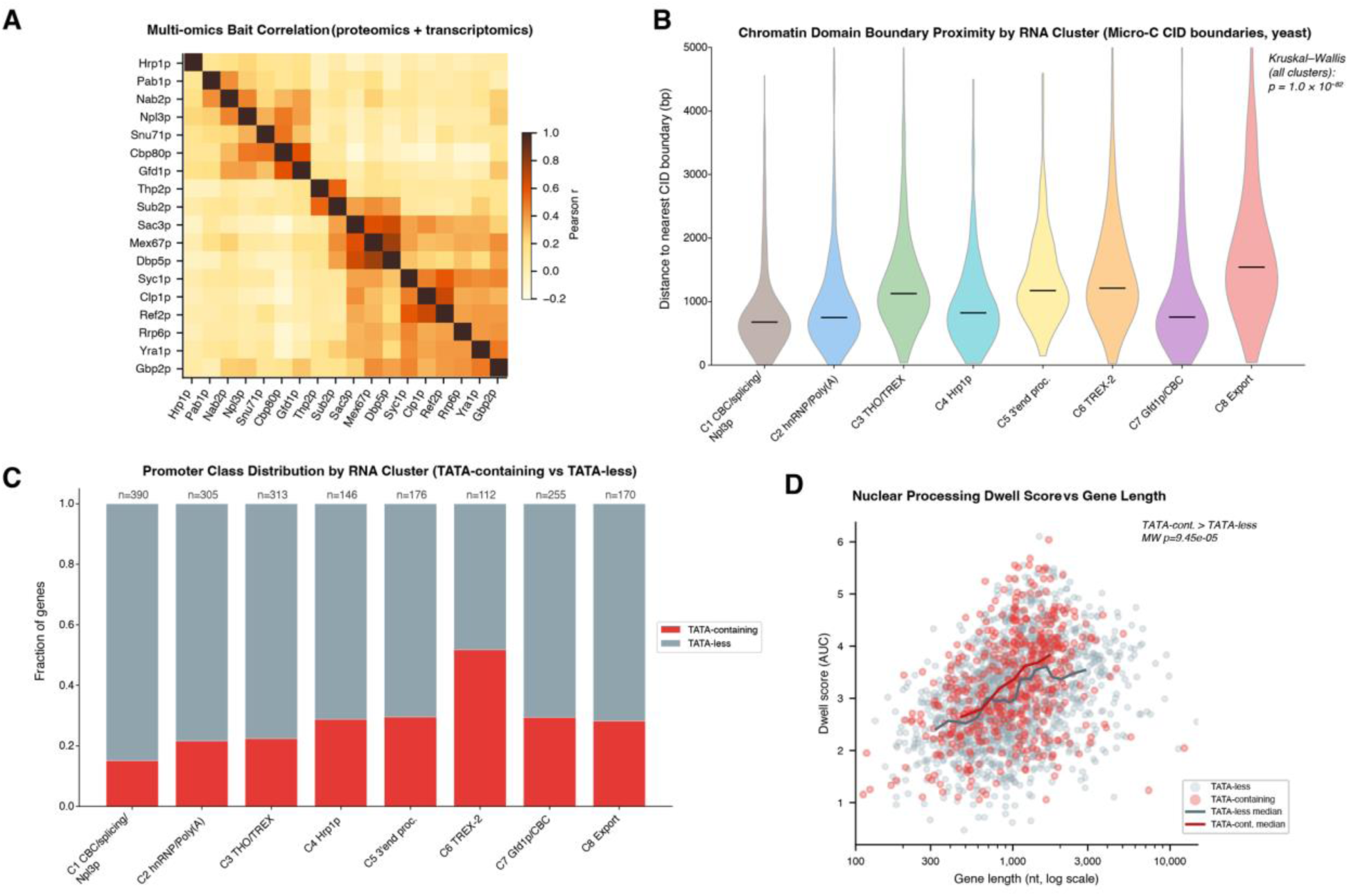
Concordant protein and RNA co-association links mRNP assembly to chromatin domains and promoter class. **(A)** Multi-omics bait–bait correlation heatmap. Combined correlation coefficients (mean of AP-MS Pearson r and AP-RNA-seq Pearson r) for all shared bait pairs, ordered by the clustering in Supplemental Figure 6A. **(B)** Chromatin domain boundary proximity by RNA cluster. Distance to the nearest Micro-C CID boundary (bp) for genes in each k = 8 transcript cluster. Violin plots; horizontal line: median. Kruskal–Wallis p = 1.0×10⁻⁸²; C1 versus each other cluster, Mann–Whitney U with Bonferroni correction, p ≤ 0.006. **(C)** Promoter class distribution across transcript clusters. Proportion of TATA-containing (SAGA-regulated) versus TATA-less (TFIID-regulated) promoters per k = 8 cluster. Promoter annotations from ref.^60^. **(D)** Nuclear processing dwell score versus transcript length, by promoter class. Dwell score (y-axis) versus ORF length (x-axis, log scale) for all co-enriched transcripts, colored by promoter class (TATA-containing: red; TATA-less: gray-blue). TATA-containing genes show elevated dwell scores across all length quartiles (Mann–Whitney U, p = 9.45×10⁻⁵).

### Chromatin domain context links mRNP co-association with gene regulatory architecture

An unexpected connection surfaced when we next asked whether the eight AP-RNA-seq transcript clusters (C1–C8; Figure 2A) were enriched for chromatin-level features. Chromosomal interaction domains (CIDs) are the structural units of yeast chromatin^55,56^, analogous to metazoan topologically associating domains (TADs)^57,58^, and their boundaries are enriched for highly expressed, SAGA-regulated genes^55^. Measuring each transcript’s distance to the nearest CID boundary revealed a striking difference: genes in the CBC/splicing/Npl3p cluster C1 lay significantly closer to CID boundaries than genes in any other cluster (Kruskal–Wallis p = 1.0×10⁻⁸²; C1 versus each other cluster, Mann–Whitney U with Bonferroni correction, p ≤ 0.006; Figure 3B). Median boundary distance ranged from 676 bp (C1) to 1,543 bp (export cluster C8), and ribosomal protein genes lay closest of all (median 683 bp versus 959 bp for non-RP genes; Mann–Whitney p = 7.5×10⁻¹⁴). Because these highly expressed, intron-containing genes are preferentially positioned near CID boundaries, this association suggests a link between the transcript classes defined by co-purification and chromatin domain architecture.

We next examined whether promoter architecture tracked with co-purification. In *S. cerevisiae*, TATA-box (SAGA-regulated) promoters drive a small set of stress-responsive, highly regulated genes, while most constitutive genes use TATA-less (TFIID-regulated) promoters^59,60^. Among the RNA clusters, the TREX-2 cluster C6 was the most enriched for TATA-containing, SAGA-regulated genes (Figure 3C; Supplemental Figure 8A). Across all transcripts, TATA-containing/SAGA-regulated genes carried a mean nuclear processing dwell score ∼0.3 units higher than TATA-less/TFIID genes (Mann–Whitney U, p = 9.45×10⁻⁵; Figure 3D; Supplemental Figure 8B), an effect that held within every gene-length quartile. Thus it is also promoter class, not length alone, that determines the breadth of biogenesis-factor association. At finer resolution, CID-boundary insulation strength showed a weak but statistically significant positive association with the mean dwell score of neighboring genes (r = 0.06, p = 1.25×10⁻³; Supplemental Figure 8C–D), a modest trend consistent with, though not on its own strong evidence for, coupling between domain insulation and local mRNP biogenesis.

Collectively, the AP-MS and AP-RNA-seq readouts of our survey defined a nuclear mRNP biogenesis network built from discrete functional modules that follow successive processing stages, linked by a small number of broadly associated factors. Among these factors, Pab1p, Yra1p and Npl3p, together with the CBC, stand out as candidate members of a common core on the basis of their broad connectivity across the network.

Ensemble purifications, however, necessarily co-isolate a mixture of mRNPs from different maturation stages and transcript identities, which limits how precisely the composition and copy number of any single mRNP can be determined. Resolving whether this organization holds at the level of a single transcript requires measuring the composition of one defined transcript directly.

### A sequential affinity purification procedure to isolate the nuclear mRNP of a single transcript

To test the survey’s observations at the level of a single, staged mRNA species, we isolated the nuclear mRNP of single defined transcripts by sequential affinity purification. We used as reporter an mCherry transcript, which is intronless, of typical yeast length (∼1 kb) and encodes a stable, biologically inert protein^61^. Because MS2 hairpin tags are a well-established handle for non-denaturing mRNP isolation in *S. cerevisiae*, we optimized the system design with the reporter transcript carrying 1, 2, 4, 6, 8 or 24 hairpins in its 3′ UTR. MS2 coat protein fused to GFP (MS2CP-GFP) was co-expressed with the MS2-tagged reporter, and captured, with the bound RNA, on anti-GFP nanobody-conjugated magnetic beads^62^. Four hairpins were sufficient for efficient recovery: co-purified mRNP proteins increased from two to four hairpins but not further with eight (Supplemental Table 5). As expected, however, a single MS2 capture recovered high amounts of cytoplasmic mRNA-binding factors (e.g., Nam7p/Upf1p and ribosomal proteins), since most cellular mRNA is cytoplasmic^63^, obscuring nuclear maturation complexes (Supplemental Figure 9A, B).

We therefore placed a nuclear-enrichment step ahead of the MS2 capture. Guided by the survey, we used well-characterized stage-defining mRNP proteins as first-step baits (Figure 1), so that each isolation sampled composition at a defined point along maturation; this sequential scheme also enriched nuclear mRNP proteins over the bulk background (Supplemental Figure 10A). To test the survey’s predicted core independently, we excluded two of the candidate core proteins, Pab1p and Yra1p, from the bait panel; by contrast, we retained the cap-binding subunit Cbp80p as the principal first-step bait, because the CBC binds the cap co-transcriptionally and is generally thought to remain bound through nuclear maturation. mCherry transcripts bearing four MS2 hairpins in their 3′ UTR (mCherry-4xMS2) were expressed from a *TDH3* promoter alongside MS2CP-GFP in strains in which the bait proteins carried a genomic PrA tag followed by an HRV-3C protease site, for elution of native mRNPs by 3C cleavage^64^ (Figure 4A). After capture of the PrA-tagged bait-associated complexes on IgG-conjugated beads, bound mRNPs were released by 3C protease cleavage, and the MS2-tagged mCherry mRNA was recaptured on anti-GFP beads, thereby isolating the nuclear mRNP assembled on that transcript at each bait’s stage.

**Figure 4.**
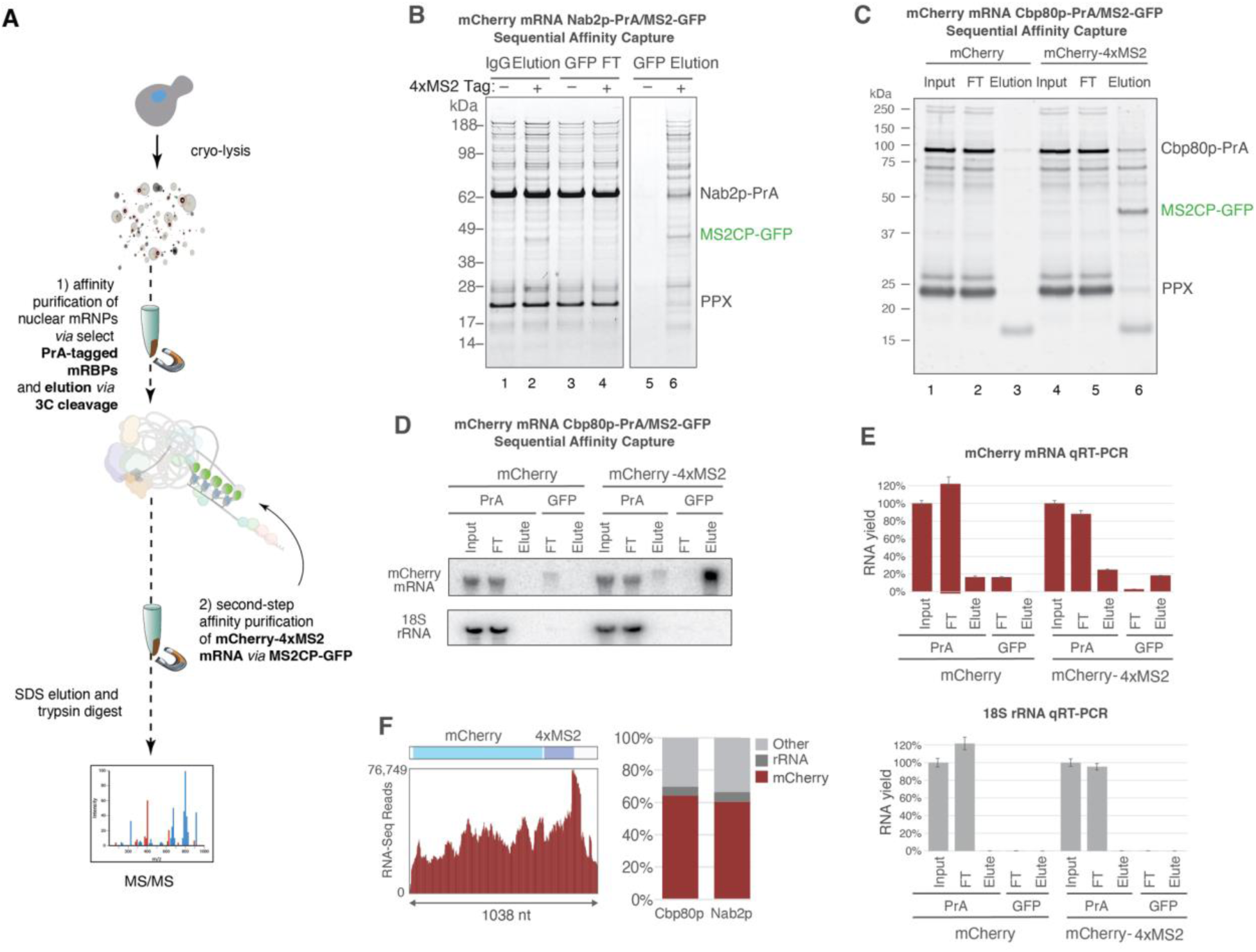
A two-step affinity scheme isolates the nuclear mRNP of a single MS2-tagged transcript. **(A)** Scheme: affinity capture of nuclear mRNPs on a PrA-tagged RNA-binding protein, on-bead HRV-3C release, then capture of the MS2-tagged mCherry mRNA on anti-GFP nanobody beads, followed by LC-MS/MS. **(B)** Coomassie-stained SDS-PAGE of the Nab2p-PrA sequential capture (IgG elution, GFP flow-through, GFP elution; ± 4xMS2 tag; lanes 1–6); MS-identified proteins marked at predicted masses. **(C)** Coomassie-stained SDS-PAGE of the Cbp80p-PrA sequential capture (input, flow-through, elution for mCherry and mCherry-4xMS2 on anti-GFP beads). **(D)** Northern blot for mCherry mRNA and 18S rRNA across purification steps. **(E)** qRT-PCR RNA yield for mCherry and 18S rRNA across steps (mean ± s.e.m., n = 3). (F) Total RNA-seq read coverage across the mCherry-4xMS2 transcript (1,038 nt) and transcript-class composition (mCherry, rRNA, other) for Cbp80p and Nab2p isolations. See also Supplemental Figure 10.

As proof of principle, we used Nab2p-PrA and Cbp80p-PrA as first-step baits, both nuclear mRNA-binding proteins that co-purified more than half of all coding transcripts in the survey (Supplemental Figure 4A) and are thus broad enough in specificity to enrich a general nuclear mRNP population^65,66^.

Sequential purification of mCherry-4xMS2 co-isolated RNP complexes in sufficient amounts to be visible by Coomassie-stained SDS-PAGE (Figure 4B, C), whereas otherwise identical transcripts lacking the 4xMS2 tag gave essentially no protein, confirming that recovery is MS2-dependent rather than through non-specific MS2CP-GFP binding (Figure 4B, lane 5; Figure 4C, lane 3; for all bait strains, Supplemental Figure 10B). In Nab2p-PrA cells the post-GFP band pattern matched the first-step eluate (Figure 4B, lanes 2 and 6), as expected for isolation of Nab2p-associated mCherry mRNPs; the decrease in Nab2p after the second step simply reflects capture of only the mCherry-bound subfraction of total Nab2p, which itself was efficiently recovered. Similar changes in the relative composition were seen in analogous sequential isolations using Cbp80p-PrA, with overall banding pattern conserved (Figure 4C, lanes 4 and 6).

Western blotting for the PrA and GFP tags across the purification confirmed high yields at each step, with MS2CP-GFP recovery strictly dependent on the 4xMS2 tag (Supplemental Figure 9C, D). RNA recovery and purity were assessed by northern blot and qRT-PCR from Cbp80p-PrA isolations (Figure 4D, E). Northern blots with mCherry-and 18S rRNA-specific probes confirmed specific recovery of full-length tagged transcript with limited degradation (Figure 4D). qRT-PCR recovered ∼20% of input mCherry mRNA after the first step (consistent with capture of only the fraction of transcripts bound by the first-step bait) and ∼75% of that after the second, with negligible 18S rRNA (Figure 4E).

Total RNA-seq provided an unbiased view of purity^67^: mCherry-4xMS2 made up ∼60% of purified RNA (Figure 4F; Supplemental Table 4). Read coverage was uniform across the full transcript, confirming isolation of intact, full-length mRNAs. Placing the tag at the 5′ end instead did not substantially change the core composition (see below).

### Copy number and mass of the Cbp80p-associated mCherry mRNP

Having established a procedure to isolate the nuclear, stage-specific pool of a single MS2-tagged transcript, we determined the protein composition of the mCherry mRNP across nuclear maturation. Co-isolated proteins were identified and quantified by label-free LC-MS/MS, and abundances were converted to copy numbers per mRNP using MS2CP-GFP as internal calibrant: each hairpin binds one MS2CP-GFP dimer, so the 4xMS2 tag carries eight MS2CP-GFP monomers per mRNP, and MS2CP-GFP signal increased with hairpin number across transcripts bearing 2, 4 or 8 hairpins (Supplemental Figure 10C). For comparison between isolations, Figures 5A and 6A show abundances relative to the Cbp80p bait.

**Figure 5.**
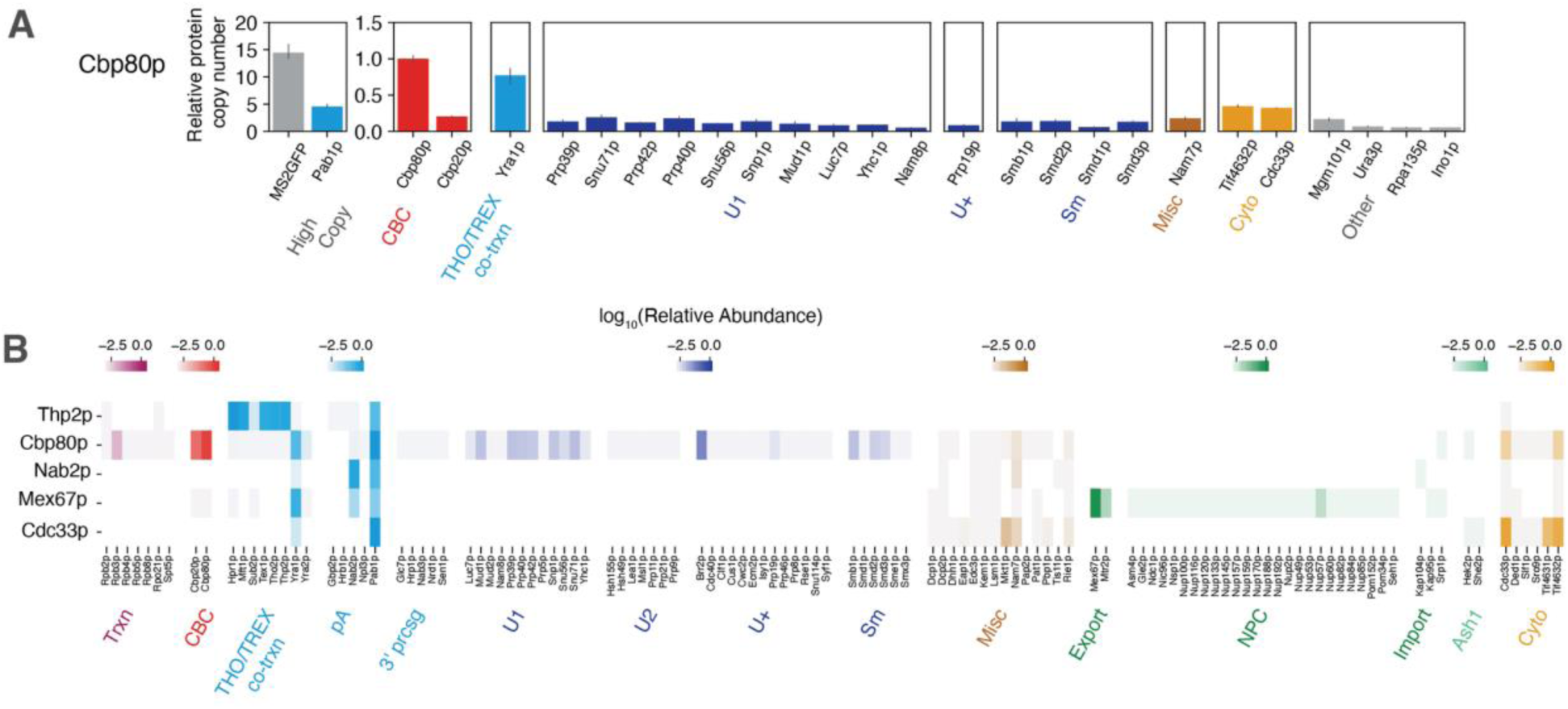
A minimal core is progressively remodeled as an mRNP matures. **(A)** Protein abundance in Cbp80p-PrA/mCherry-4xMS2 mRNPs relative to Cbp80p (set to 1); per-mRNP copy numbers calibrated to MS2CP-GFP (eight per mRNP) are given in Supplemental Table 6; high-copy proteins are shown on a separate scale, and proteins below 5% of Cbp80p are omitted. The glutaraldehyde-stabilized version of this quantification (3 mM) is shown in Supplemental Figure 10F, with a 0–3 mM titration for representative proteins in S10E. **(B)** Protein-abundance heatmap across five stage-defining baits—Cbp80p (CBC), Thp2p (co-transcriptional), Nab2p (nuclear poly(A)), Mex67p (export) and Cdc33p (cytoplasmic)—normalized to each tagged bait; proteins below 1% in all samples are omitted. See also Supplemental Figure 9.

**Figure 6.**
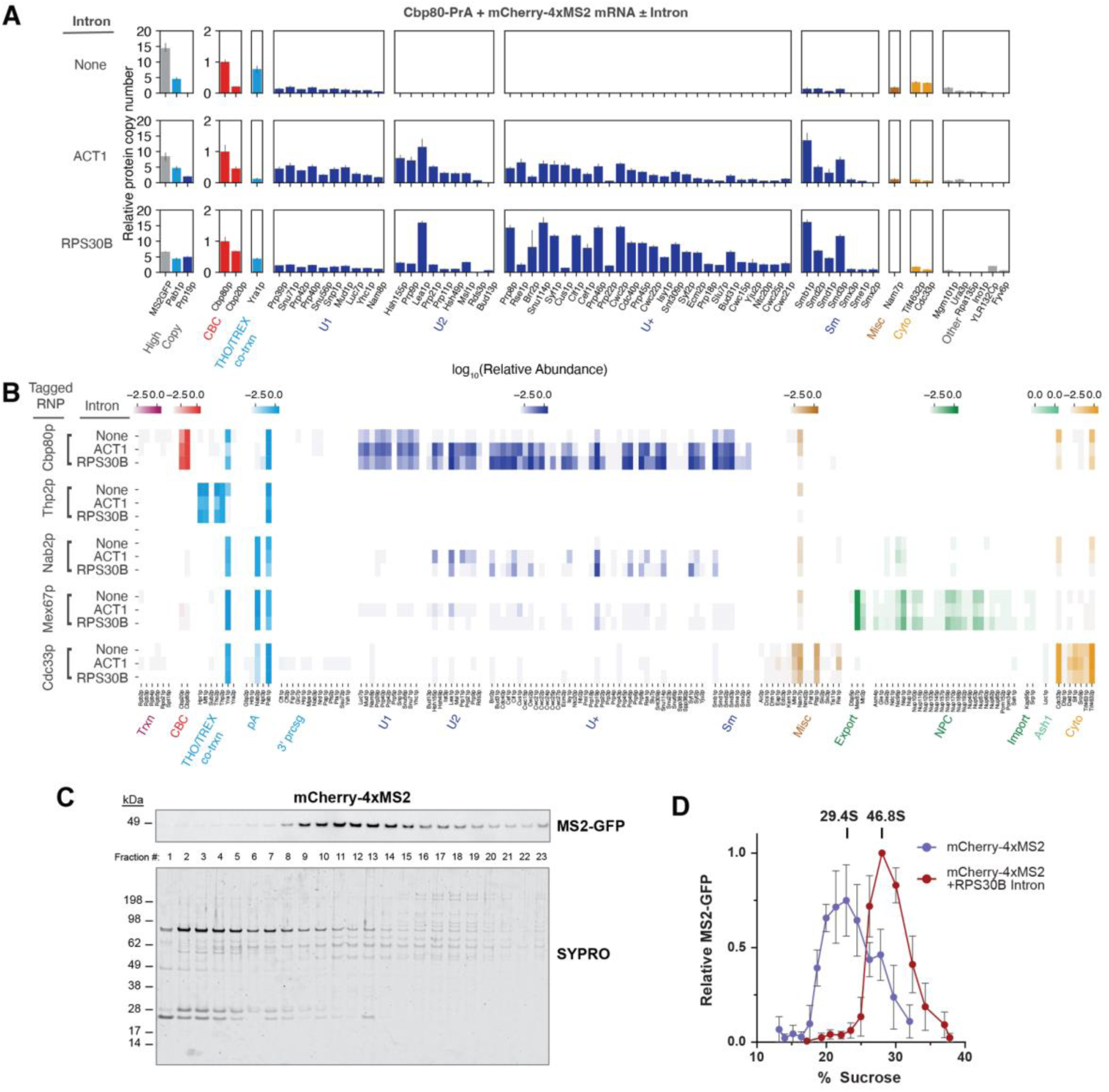
**Intron class, not intron presence** alone, determines spliceosome composition and mRNP size. **(A)** Relative protein copy number for Cbp80p-PrA/mCherry-4xMS2 mRNPs with no intron, the *ACT1* intron, or the *RPS30B* intron, normalized to Cbp80p; high-copy proteins on a separate scale. **(B)** Protein-abundance heatmap of the intron variants across stage-defining baits, normalized to each tagged bait. **(C)** Sucrose-gradient sedimentation of Cbp80p-PrA-purified mRNPs: representative SYPRO-stained gel and MS2CP-GFP western blot across fractions. **(D)** MS2CP-GFP sedimentation profile for mCherry-4xMS2 with or without the *RPS30B* intron (peaks at 29.4S and 46.8S; mean ± s.e.m., n = 3). See also Supplemental Figure 9.

Quantitative MS of Cbp80p-PrA-associated mCherry-4xMS2 mRNPs returned strikingly few abundant proteins (Figure 5A). Per mRNP, Pab1p was present at ∼2.5 copies, consistent with occupancy upstream of the poly(A) site and along the poly(A) tail, and Yra1p, recruited co-transcriptionally and released before export, at ∼0.4–0.5 copies on average, with and without short in-lysate glutaraldehyde stabilization^17,40,68–70^. The CBC subunits were the only core components that were recovered at lower levels without (Cbp80p ∼0.55, Cbp20p ∼0.1 copies) compared to with glutaraldehyde stabilization (∼1.1 and ∼0.6 copies; Supplemental Table 6), suggesting that the CBC, and Cbp20p in particular, can be more dynamically associated with the mRNP than generally assumed. All other maturation factors—THO/TREX subunits, spliceosomal proteins, Npl3p and Nab2p—were present at sub-stoichiometric levels (Figure 5A; Supplemental Table 5), as expected for dynamic, transient association across a Cbp80p-bound population that spans every stage from early nuclear assembly to export; the U1 snRNP components among them are consistent with promiscuous splice-site scanning, in which U1 transiently surveys nascent transcripts before committing to bona fide 5′ splice sites^71^. Only the CBC, Yra1p and Pab1p were present at high copy number. This is precisely the core highlighted by the mRNP ensemble survey, where the same factors emerged as the most broadly connected nodes of the co-purification network; the single-mRNP measurement established them as both broadly associated and consistently present, at defined copy numbers, on an individual transcript.

Moreover, a control experiment using Cbp80p-PrA-purified mRNPs mixed with heavy-lysine-labeled wild-type lysate determined heavy-to-light ratios by MS to confirm that measured mRNP composition was not distorted by post-lysis exchange^72^. CBC subunits and the MS2CP-GFP calibrant were stable over one hour (<5% exchange) and Pab1p exchanged up to ∼30%, without affecting its copy number per mRNP, which matched that measured with and without glutaraldehyde stabilization (Supplemental Figure 10D–F). Together with the recovery of the expected stage-specific complexes and core proteins (Figure 5), this indicates that the sequential purification captures bona fide mRNPs matured through the same pathway as endogenous mRNAs.

From the RNA sequence and MS-quantified protein abundances of a glutaraldehyde-stabilized complex we estimated an average mass of ∼1.2 MDa per mCherry-4xMS2 mRNP, or ∼0.8 MDa without the MS2CP-GFP tag and its hairpins (Supplemental Figure 10F; Supplemental Table 6). Of its ∼540 kDa of protein on ∼280 kDa of RNA, about half (∼290 kDa) is contributed by the core components, Yra1p, Pab1p and the more dynamically associated CBC (Cbp80p, Cbp20p), and the rest by transiently associated factors, each present on only a fraction of particles. Protein thus accounts for the majority of the particle’s mass (∼2:1 protein:RNA after glutaraldehyde stabilization, ∼1.3:1 without). Together with its compact sedimentation behavior (∼29S; see below), this is consistent with the compact nuclear mRNPs described by cryo-electron tomography^29^.

### Composition of a single mRNP across successive maturation stages

To follow composition along the pathway rather than from a single vantage point, we sequentially isolated mCherry-4xMS2 complexes from additional stage-defining baits: Thp2p, a THO/TREX subunit recruited co-transcriptionally to elongating RNA polymerase II; Nab2p, a general nuclear poly(A)-binding protein; Mex67p, the principal mRNA export receptor; and Cdc33p (eIF4E), which replaces the CBC on the cap^1,3^. MS2-dependent co-isolation was confirmed for each bait by Coomassie-stained SDS-PAGE (Supplemental Figure 10B).

Protein compositions shifted markedly with the stage of maturation sampled but always contained the same small core subset (Yra1p and Pab1p; Figure 5B). Beyond the core, Thp2p-associated mRNPs were enriched for THO subunits (Hpr1p, Mft1p, Tho2p) and the TREX helicase Sub2p, capturing the early co-transcriptional stage at which the export-adaptor machinery is actively loaded. Nab2p-associated isolations returned a near-minimal mRNP of Yra1p and Pab1p alongside Nab2p, as expected for a polyadenylated transcript carrying only core factors. Mex67p-associated mRNPs were enriched for its obligate partner Mtr2p and for Yra1p (the adaptor that recruits Mex67p), with Pab1p, Nab2p and low but consistent levels of NPC components present, marking an export-competent particle; Cdc33p-associated mRNPs had exchanged the nuclear cap-binding complex for the cytoplasmic translation-initiation machinery, marking the transition toward a cytoplasmic mRNP.

Across all five baits, then, composition was highly dynamic: processing complexes come and go, while Yra1p and Pab1p persist with every bait from the earliest nuclear stage through export, and the CBC with the cap-bound mRNP until its exchange for Cdc33p. These isolations thus directly confirm the core highlighted by the mRNP survey and show that THO/TREX, spliceosomal and export-adaptor modules load onto that core dynamically rather than constitutively.

### Intron class, not presence alone, determines spliceosome composition and mRNP size

In our survey, intron-containing transcripts were concentrated in two clusters, with RP and non-RP intron genes segregating into C1 and C7, respectively; we therefore asked how an intron changes the composition of a single mRNP. We inserted the well-characterized *ACT1* intron (309 nt, 31 bp into the mCherry coding sequence)^73^ into mCherry-4xMS2 and isolated it with Cbp80p-PrA as first-step bait, as the CBC binds to the capped mRNA prior to splicing and promotes early spliceosome assembly^74,75^. The intron-containing mRNP recovered all major spliceosomal subcomplexes, in contrast to the intronless transcript, which yielded only the sub-stoichiometric subset of U1 snRNP components seen before (Figure 6A); the intron-containing construct also yielded more mRNP (Figure 6A), consistent with intron-mediated enhancement of gene expression^76,77^. Critically, the ratio of Pab1p to the CBC was unchanged by the intron, and Yra1p remained present, albeit at lower levels (Figure 6A; Supplemental Table 5); splicing thus adds a transient module while the core components remain.

To test whether the survey’s RP/non-RP split reflects intron identity rather than length, we compared two similarly sized introns of the two classes in the mCherry reporter transcript: the non-RP *ACT1* intron (309 nt) and the RP *RPS30B* intron (411 nt). Isolated by Cbp80p-PrA sequential purification to maximize spliceosome yield, the two gave distinct subcomplex compositions (Figure 6A, B). U1 and U2 snRNPs, comprising the early prespliceosome (complex A), were enriched ∼2–3-fold on *ACT1*-intron containing mCherry mRNAs, whereas the activated (Bact) and catalytic (C) spliceosome, including the entire Prp19 complex, were enriched ∼2–3-fold on *RPS30B*-intron expressing mCherry transcripts. Because the two introns have similar lengths, these differential enrichments reflect intron class, not length—the same division we observed in the survey between the RP (cluster C1) and non-RP (cluster C7) intron populations, i.e., it appears that the ACT1 intron accumulates at the prespliceosome while the *RPS30B* intron commits more efficiently to catalysis, consistent with the distinct splicing behavior of RP pre-mRNAs in spliceosomal mutants^78^.

Commitment to catalysis was also supported by particle size: Cbp80p-PrA eluates from cells expressing mCherry-4xMS2 were resolved on sucrose gradients and the reporter mRNP followed by MS2CP-GFP western blotting across fractions. The intronless mRNP peaked at ∼29S (∼0.8–1 MDa^79^; the MS estimate for the MS2CP-GFP-bound particle is ∼1.2 MDa), whereas the *RPS30B*-intron transcript sedimented markedly faster (∼47S, ∼2 MDa), consistent with a stably associated spliceosome (∼1.6 MDa in its activated form^80^) and with the 40–45S peaks reported for other yeast spliceosome isolations^81^ (Figure 6C, D). As with the *ACT1* intron, the core was retained: the Pab1p:CBC ratio was unchanged and Yra1p was again present (Figure 6A; Supplemental Table 5).

In addition, an observation from the mRNP survey was borne out at single-transcript resolution. Npl3p was co-isolated only with the *RPS30B* (RP) intron, not with the *ACT1*-intron-containing or intronless transcripts (Figure 6B), in line with our mRNP survey, where Npl3p preferentially co-purified RP intron-containing mRNAs (cluster C1; roughly twofold over non-RP intron and intronless transcripts). This is also consistent with its SR-like role in promoting splicing of a subset of intron-containing genes^15,82,83^. The intron-specific recruitment seen here on a single transcript is thus consistent with what was observed in our mRNP survey data, where RP pre-mRNAs clustered together in C1, apart from the non-RP intron cluster C7, with preferential Npl3p association.

### 3′-UTR elements add dedicated factors to an invariant core

Finally, we asked which transcript features change mRNP composition and which leave the core untouched, questions the survey could address only in a correlational manner. We compared mCherry-4xMS2 transcripts differing in a defined 3′-UTR element, promoter or length, as well as a transcript in which the mCherry coding sequence was replaced by that of *URA3* (Figure 7A).

**Figure 7.**
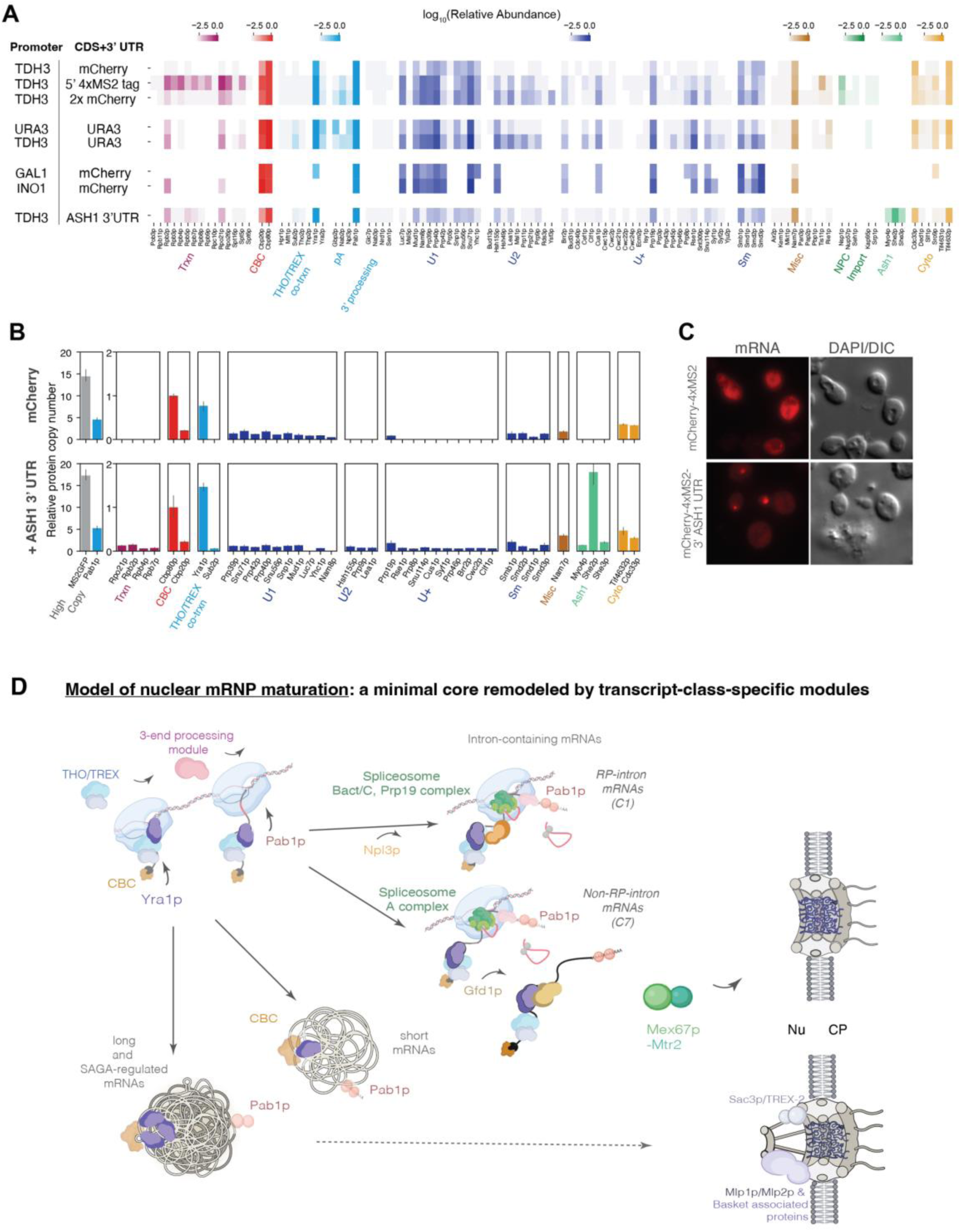
Core composition is robust to sequence, promoter and length, while 3′-UTR elements recruit specific factors. **(A)** Protein-abundance heatmap for mCherry-4xMS2 variants—5′-placed tag, tandem 2×mCherry, *URA3* or mCherry from endogenous or *TDH3* promoters, *GAL1*-and *INO1*-driven transcripts, and the *ASH1* 3′-UTR construct—by Cbp80p-PrA capture, grouped by processing role. **(B)** Relative protein copy number with or without the *ASH1* 3′-UTR, normalized to Cbp80p; high-copy proteins on a separate scale. **(C)** Single-molecule FISH of mCherry-4xMS2 and mCherry-4xMS2-*ASH1*-3′UTR transcripts (mRNA channel and DIC); the *ASH1*-tagged transcript localizes to the bud tip. **(D)** Model of nuclear mRNP maturation: a minimal core remodeled by transcript-class-specific modules, integrating our findings with established steps of mRNP biogenesis. Co-transcriptionally, Yra1p and Pab1p, together with the more dynamically associated CBC, form a minimal core onto which THO/TREX and 3′-end processing modules associate transiently (curved arrows). On RP-intron mRNAs (cluster C1), which associate with Npl3p, the spliceosome progresses to the activated/catalytic stage (Bact/C, Prp19 complex), whereas on non-RP-intron mRNAs (cluster C7) it accumulates at the A complex, and these transcripts subsequently associate with Gfd1p. mRNAs are exported by the Mex67p–Mtr2p heterodimer, short and intron-containing mRNAs mainly through basket-less NPCs, and long and SAGA-regulated mRNAs through NPCs carrying TREX-2 (Sac3p) and the nuclear basket (Mlp1p/Mlp2p). Nu, nucleus; CP, cytoplasm.

We appended the *ASH1* 3′ UTR, whose stem-loops recruit She2p and She3p to carry the transcript to the bud tip along actin filaments via the myosin Myo4p^84–87^, to the *TDH3*-driven mCherry-4xMS2 construct and isolated it with Cbp80p-PrA (Figure 7B). Fluorescent in situ hybridization (FISH) microscopy using a probe against mCherry mRNA confirmed bud-tip localization of the transcript, as for native *ASH1* mRNA (Figure 7C), and MS recovered abundant She2p, She3p and Myo4p, showing that the isolation captures element-specific factors with high sensitivity and specificity, further validating the single-transcript approach^88^. Loc1p and Puf6p, reported to associate with *ASH1* mRNA in the nucleus, were absent, consistent with their more transient association and displacement during export^89^. A defined 3′-UTR element thus adds its own dedicated factors on top of the otherwise invariant core.

In contrast, the survey had linked promoter class to the breadth of biogenesis-factor association across the transcriptome (Figure 3D), so we asked whether promoter identity also reshapes the core of an individual mRNP. Compared with mCherry expressed from the *TDH3* promoter, mCherry driven by the inducible SAGA-regulated INO1 or *GAL1* promoters, *URA3* expressed from either its endogenous or the *TDH3* promoter, and a tandem 2×mCherry transcript (doubling transcript length and sequence)^90^ all showed unchanged core composition and copy number after isolation via Cbp80p-PrA (Figure 7A)^59,90^. Only the SR-like proteins Gbp2p and Npl3p differed, being modestly more abundant on the *URA3* transcripts than on the single mCherry transcript, with the 2×mCherry transcript intermediate. These shuttling SR-like proteins act in surveillance and 3′-end formation^91–93^ and may recognize sequence features of the transcript beyond their known preference for intron-containing mRNAs^15,94,95^. A mCherry transcript carrying the 4xMS2 array at its 5′ end instead likewise gave a similar core profile, except for markedly higher recovery of RNA polymerase II subunits and, to a lesser extent, U1 snRNP proteins, consistent with capture of nascent, co-transcriptional mRNPs. Taken together, the core measured in Cbp80p-associated mRNPs—the CBC, Yra1p and Pab1p—proved essentially invariant to promoter, sequence or length, providing single-transcript evidence that the promoter-class effect seen in the survey acted on the breadth of factor association, not on the minimal core.

## DISCUSSION

### A minimal shared core and staged, modular assembly of mRNPs

The composition and dynamics of nuclear mRNPs have been difficult to define, because, with ensemble data alone, a small set of universal components cannot be unambiguously distinguished from the much larger set of RBPs that associate transiently during processing. By pairing two genome-scale approaches—an affinity-purification survey of 34 mRNP-associated proteins profiled by proteomics, 18 of them also by transcriptomics of the same isolations, and a sequential single-transcript purification that resolves composition stage by stage—we distinguish the common and transient components. This combination is essential: ensemble co-association cannot on its own distinguish a stably bound factor from one sampled broadly but dynamically, and single-transcript proteomics of one reporter cannot establish generality; only a factor identified as broadly associated across the survey and then found consistently present on individual reporter transcripts through successive stages can be assigned to the core. On this basis, the co-transcriptionally recruited export adaptor Yra1p and the poly(A)-binding protein Pab1p, together with the more dynamically associated cap-binding complex (Cbp80p/Cbp20p), form a minimal nuclear core that persists from co-transcriptional assembly to export, while all other factors, such as THO/TREX, spliceosomal subunits, cleavage and polyadenylation factors, and the SR-like proteins Npl3p and Gbp2p, associate sub-stoichiometrically and in a stage- or transcript-specific manner. The resulting particle is compact (∼0.8 MDa untagged, ∼1.2 MDa with the MS2CP-GFP reporter, ∼29S, with protein accounting for the majority of its mass), and its core is unaltered by promoter, transcript length or coding sequence.

That so few proteins suffice to specify a maturing particle argues that nuclear mRNP biogenesis in yeast follows a minimalist, staged assembly and exchange rather than the continuous accumulation of a large dedicated assembly; the mRNP thus behaves as a modular scaffold that exchanges compositionally distinct RBP assemblies at successive biogenesis checkpoints, the core providing continuity and the stage-specific modules conferring functional identity. The timing of this modular loading is itself informative: within the export machinery, Thp2p is recruited co-transcriptionally as part of THO, while Sub2p is subsequently transferred to the nascent RNA^54^, and Pab1p engages early at the poly(A) site, indicating that the order of module recruitment, not module identity alone, shapes mRNP maturation.

Yra1p’s centrality in both datasets concurs with and extends recent structural and single-molecule work. Cryo-electron tomography and crosslinking mass spectrometry depicted nuclear mRNPs as compact particles in which the positively charged disordered regions of Yra1p or its human ortholog ALYREF drive the RNA–RNA interactions that compact the transcript, a packaging function conserved from yeast to metazoans^29^. Our findings are consistent with such Yra1p-mediated packaging imposing a default, sequence-independent architecture onto which other factors load transiently. Single-molecule imaging likewise reported Yra1p on individual transcripts with pronounced molecule-to-molecule heterogeneity^25^. Our single-mRNP measurements give an average of ∼0.4–0.5 Yra1p per mRNP. Because the Cbp80p-bound population spans every stage up to export, and Yra1p is released from the mRNP before export^1^, this average likely reflects both the molecule-to-molecule heterogeneity seen by single-molecule imaging^25^ and the removal of Yra1p at late stages. Yra1p is thus a core component in that it accompanies the mRNP throughout maturation, rather than being bound to every particle at every moment.

The same study^29^ also identified Yhs7p as a component of these nuclear mRNPs, in close proximity to Yra1p. Yra1p–Yhs7p crosslinks, together with AlphaFold-predicted interfaces between the RRM domain of each protein and a short helical segment of the other, placed the two proteins adjacent to one another. In addition, *in vitro* RNA-annealing assays showed that Yhs7p, like Yra1p, promotes RNA–RNA interactions through its positively charged disordered regions^29^. Together, these observations define a yeast Yra1p–Yhs7p module analogous to the human Aly/REF–ERH–CHTOP module^29^. Consistent with an early mRNP biogenesis role close to Yra1p, the Yhs7p bait co-purified Yra1p in our survey, and live imaging has placed it among the earliest mRNP proteins at transcription sites, co-arriving with Yra1p and Cbp80p^96^.

Normalized copy numbers showed that the CBC was the only core component whose recovery rose with glutaraldehyde stabilization, most strongly for Cbp20p (Supplemental Figure 10E, F). Although the small size of Cbp20p (∼24 kDa, few detectable peptides) may lower its signal, the stabilization-dependent increase suggests that the CBC, and possibly the Cbp80p–Cbp20p heterodimer itself, associates with the mRNP more dynamically than generally assumed; with glutaraldehyde stabilization, CBC recovery approached one copy per mRNP, consistent with each particle carrying a CBC.

Overall, our results place earlier surveys of yeast mRNP composition on a firmer mechanistic footing. Transcriptome-wide crosslinking had shown that Nab2p, Npl3p and Yra1p bind broadly across coding regions, with occupancy modulated by features such as intron content^17^, and that apparent mRNP composition varies with transcript sequence^16^. By resolving how many of those contacts persist on a single mRNP, we show that such broad binding reflects dynamic, sub-stoichiometric sampling rather than stable occupancy, and that the three-protein core is itself sequence-independent; the class-to-class differences in mRNP complexity reported previously^13,16,17^ thus arise mainly from the diversity and stage specificity of transiently associating processing factors, rather than variation in the stable core. What the combination of survey breadth and single-transcript resolution adds, beyond either alone, is direct evidence that a maturing mRNA carries a small, defined, sequence-invariant core throughout maturation, while its variable proteome is the dynamic, transcript-class-specific machinery layered upon it.

### A conserved core with a metazoan-specific link between splicing and export

The core and its assembly appear broadly conserved. Its three components have clear metazoan counterparts—the CBC (NCBP1/NCBP2), nuclear and cytoplasmic PABP, and the Yra1p homolog ALYREF—all acting in packaging and export, and cryo-EM of the human THO–UAP56 complex showed that UAP56/DDX39B couples ATP turnover to ALYREF loading on TREX and its handover to the TREX-2/NPC system^30^. Here, Sub2p, the yeast UAP56 ortholog, co-purified with Yra1p in Thp2p-associated co-transcriptional mRNPs. This marks the loading step as ancient (at least common to the opisthokont branch of eukaryotes) and builds on foundational yeast biochemical work in which co-transcriptional Sub2p recruitment to THO, followed by Yra1p loading, licenses export^39,97,98^.

Where yeast and metazoa diverge is in how the completion of splicing is recorded. Metazoans deposit the exon junction complex (EJC) at exon–exon junctions, coupling splicing to export and surveillance^99^, and cryo-EM has shown each EJC to serve as a docking platform for ALYREF that recruits TREX through multivalent contacts, compacting the mRNP into a periodic assembly^28^—a permanent, one-per-junction record of splicing. Yeast has no EJC, and consistent with this our intron-containing isolations carry no comparable splicing-deposited mark: the intron adds only a transient spliceosomal module and leaves no permanent, per-junction record. Its ALYREF ortholog Yra1p is not deposited at junctions but is loaded co-transcriptionally as part of the core, present on a subset of mRNPs at any one time, with no consistent change in its average occupancy when an intron is present (Figure 6A). Thus in yeast, export licensing is uncoupled from any record of splicing, whereas in metazoans ALYREF docking is EJC- and splicing-coupled. The modular organization of a core scaffold that exchanges stage-specific assemblies at successive checkpoints is therefore conserved from yeast to human, although the means of recording checkpoint completion is not, being transient in yeast and permanently inscribed by EJC deposition in metazoa. Consistent with a lighter regulatory layer in yeast, the SR-like proteins Npl3p and Gbp2p associate only conditionally in our data (with intron-containing or *URA3* transcripts), in contrast to the broad role of metazoan SR proteins in splicing and export.

### Npl3p, intron class and coupling to genome organization

Npl3p illustrates how our survey and single-mRNP analyses complemented one another and refined our understanding of nuclear mRNA maturation. In our mRNP ensemble survey, Npl3p emerged as one of the most broadly connected nodes in the mRNP network, alongside Pab1p and Yra1p, as a candidate core component. Yet in the single-mRNP analysis it was present only sub-stoichiometrically in Cbp80p-associated mRNPs and was recruited specifically by the *RPS30B* intron, not by the *ACT1* intron or the intronless transcripts.

Importantly, the two approaches sampled different transcript populations. The survey interrogated the whole endogenous transcriptome, in which many highly expressed mRNAs—RP mRNAs above all—carry introns and other features that recruit Npl3p; its high network connectivity thus reflected broad but conditional association, concentrated in the CBC/splicing/Npl3p cluster C1. Our single-mRNP analysis of a non-endogenous intronless model transcript isolated the minimal, sequence-independent core, of which Npl3p was not a constitutive part, consistent with recent single-molecule data, which likewise found substantial Npl3p heterogeneity across individual mRNP molecules^25^. These observations make Npl3p a conditional component; broadly associated with the endogenous mRNP proteome because its preferred substrates are abundant, but not part of the minimal, sequence-independent scaffold. Mechanistically, this rests on Npl3p’s role as the yeast SR-like splicing factor which promotes U1 and U2 snRNP recruitment to a broad cohort of intron-containing genes^15^, the molecular basis for our observed *RPS30B*-specific enrichment. Its phosphorylation state, set by the Sky1p kinase and Glc7p phosphatase and paralleling the dephosphorylation-dependent export of mammalian SR proteins^9,100,101^, further couples its mRNP association to export competence.

Our analysis of intron-type-specific remodeling exposes a previously underappreciated layer of co-transcriptional regulation. The contrasting enrichment of pre-spliceosomal U1/U2 snRNPs with the *ACT1* intron versus the activated Bact and catalytic C spliceosome, including the entire Prp19 complex, with the *RPS30B* intron identified distinct kinetic bottlenecks for transcripts encoding these different intron classes. RP-gene introns like *RPS30B* appeared to commit more efficiently to catalysis, consistent with the distinct behavior of RP transcripts in spliceosomal mutants^78^. This was mirrored transcriptome-wide in the mRNP survey, where intron-containing genes, RP genes in particular, clustered together (C1) and were strongly enriched for Npl3p co-purification. Given this role in U1 and U2 snRNP recruitment^15^, Npl3p’s preferential association with RP pre-mRNAs may contribute to their faster progression to catalytic activation.

Notably, these intron-containing RP genes also lay closest to chromosomal interaction domain (CID) boundaries. Why the most highly transcribed, co-regulated RP genes should lie closest to domain boundaries is unclear. Their shared position may reflect a higher-order spatial organization of the genome that places co-regulated transcripts with similar processing requirements near one another, coupling mRNP assembly to chromatin architecture, though it may equally follow from the high transcriptional activity at which CID boundaries themselves form.

### TREX-2 and the nuclear basket define a selective export checkpoint

TREX-2 (Sac3p/Thp1p/Sus1p), which links the SAGA transcriptional co-activator to the mRNA export machinery^102–104^, has recently been shown by cryo-electron tomography to be an integrally associated module of the NPC in vertebrates^44^ and in yeast (Soni *et al.*, manuscript in preparation). Our data reinforce this notion and extend it to the transcript level: the TREX-2 bait Sac3p co-purified nucleoporins as well as the Mlp1p/Mlp2p basket proteins, placing TREX-2 at basket-containing NPCs^2^. The transcripts most enriched with Sac3p formed a discrete cluster (C6) of SAGA-regulated, stress-inducible genes, these being classic targets of gene recruitment to the NPC by Sus1p–TREX-2^102,103,105,106^. This enrichment was specific to Sac3p and not shared by other export-related baits, such as Mex67p or Dbp5p. Besides SAGA-regulated and stress-inducible genes, higher Sac3p association was also observed with longer transcripts (clusters C5 and C8), in line with the preferential export of longer mRNAs through TREX-2 and basket-containing NPCs^2^.

Direct RNA-level evidence for a basket checkpoint comes from cross-correlation with transcripts associated with basket-containing NPCs in the Mlp1p AP-RNA-seq dataset of Bensidoun *et al*.^2^, in which RNAs co-purifying with Mlp1p, a principal scaffold of the NPC’s nuclear basket, were compared between basket-containing and basket-less NPC affinity-purified samples. Mlp1p RNA profiles correlated most strongly with the RNA profiles from late-export/TREX-2 baits of our survey (Spearman ρ: Dbp5p 0.40, Sac3p 0.38, Clp1p 0.35), and transcripts co-purified by these baits were those preferentially enriched at basket-containing over basket-less NPCs (Sac3p ρ = 0.48, Clp1p 0.44, Dbp5p 0.37 with the basket-containing/basket-less log2 fold change). The TREX-2 cluster (C6) was the most enriched at basket-containing NPCs (median log2 fold change +0.97; Mann–Whitney U vs. all other clusters, p = 1.4×10⁻⁴⁸), followed by the export cluster C8, while splicing-stage transcripts (C1) were the most depleted (−1.01; p = 1.2×10⁻⁷²), consistent with their export mainly through basket-less NPCs. These Mlp1p data support a model in which the nuclear basket is assembled on a subset of NPCs and acts as a quality-control checkpoint for export-directed mRNPs, within which TREX-2 confers additional selectivity for the SAGA-regulated subset. Together, these observations suggest that genome spatial organization, transcriptional regulation, mRNP assembly and NPC quality control are likely coupled^2^.

How mRNPs form on transcripts of differing lengths, and hence vary widely in size, and how they are accommodated by the size-limited central channel of the NPC for their export remain open questions^107^. However, the preferential engagement of longer transcripts by TREX-2 and nuclear basket-containing NPCs points to length-dependent organization at the export interface, although the underlying particle architecture, for example whether the mRNPs of long transcripts are arranged as strings of repeating channel-compatible subparticles as proposed for Balbiani-ring mRNPs^108,109^, will require direct structural analysis.

Taken together, nuclear mRNP maturation is neither a single universal pathway acting identically on every transcript nor a fully transcript-tailored one, but instead a hybrid: a sequence-invariant core accompanies the mRNP through maturation and is remodeled by stage-specific modules that exchange as pre-assembled units rather than as individual proteins, and this remodeling is tuned by intron class, promoter class and the spatial organization of the genome (Figure 7D). The core proteins (CBC, Yra1p, Pab1p) can be read as a minimal ‘mRNPome’—the protein content sufficient to define a nuclear transcript during maturation—while the far larger surveyed proteome is the dynamic, transcript-class-specific machinery that elaborates this core co-transcriptionally. Notably, this organization parallels ribosome biogenesis, where a stable RNA–protein core assembles co-transcriptionally while many biogenesis factors associate only transiently^110^, suggesting a general principle for large RNA–protein assembly pathways. With the core conserved from yeast to human, the added complexity of metazoan mRNPs—EJC deposition, an expanded SR-protein repertoire, pervasive alternative splicing—likely represents elaborations on this conserved core, and the combined multi-omics and single-mRNP framework applied here offers a template for dissecting mRNP biogenesis at comparable resolution in other systems.

### Limitations of the study

Some limitations bear on interpretation. First, although the sequential-isolation approach applies to any MS2-taggable transcript, we defined the minimal core primarily on one non-endogenous model transcript and compared only two introns (*ACT1* and *RPS30B*). Core composition was invariant across a panel of sequence, promoter and length variants, but endogenous mRNAs—particularly those with complex secondary structures, regulatory elements (as shown for *ASH1*) or alternative polyadenylation sites—may carry additional compositional features, and two introns cannot span the diversity of intron-dependent remodeling, particularly in metazoans, where >90% of genes carry introns compared with <5% in yeast. The generality of the minimal core and of our intron-kinetics framework therefore awaits confirmation on further endogenous transcripts and introns. Nevertheless, replacing the mCherry coding sequence with that of an endogenous gene (*URA3*), expressed from its own or the *TDH3* promoter, left the core unchanged (Figure 7A). Second, sequential affinity capture provides ensemble snapshots at bait-defined stages; it does not resolve the real-time kinetics of individual assembly events or necessarily capture rare, transient intermediates. Third, despite 34 bait proteins, the survey is a curated subset of the yeast mRNP proteome, and additional low-abundance or redundant RBPs may contribute in ways not captured here. Fourth, the CID-boundary and promoter-class analyses are correlative; direct perturbation of chromatin-domain architecture will be needed to test whether CID positioning actively influences processing efficiency. Fifth, glutaraldehyde stabilization affects what is recovered. The AP-RNA-seq purifications were not stabilized and therefore report steady-state occupancy weighted by nuclear dwell time, so the enrichment of longer transcripts in the late-export clusters (notably C8) likely reflects both residence time and affinity; nevertheless, bait-specific signals defined by gene class, such as the TREX-2 preference for the SAGA regulon, not shared by Mex67p or Dbp5p, remain robust. Likewise, per-mRNP copy numbers are calibrated to MS2CP-GFP, assuming full occupancy of the four hairpins, and CBC recovery depended on glutaraldehyde stabilization, so CBC copy numbers from unstabilized isolations are likely underestimates. Within these limits, our results define a minimal core of Yra1p and Pab1p, together with the more dynamically associated CBC, on the transcripts tested; identify Npl3p as a conditional, intron-class-dependent component; show that intron class sets spliceosome composition; and link mRNP assembly to promoter class and to the positioning of genes near CID boundaries.

## RESOURCE AVAILABILITY

New data reported in this article are available in the Supplemental Tables. Mass spectrometry data will be deposited in the ProteomeXchange Consortium via the PRIDE partner repository, and RNA-sequencing data in the Gene Expression Omnibus (GEO). Accession numbers will be made available upon deposition. Analysis code is available at GitHub (https://github.com/oeffingerlab/mRNP-survey-yeast).

## SUPPLEMENTAL INFORMATION

Supplemental Figures 1–10 and Supplemental Tables 1–6.

### Supplemental Tables

**Supplemental Table 1.** AP-MS proteomics data. (A) Raw spectral counts per bait–prey pair (LC-MS/MS survey). (B) Curated total-spectral-count matrix (proteins × pull-down runs, WT single-step APs). (C) Per-bait-normalized prey/bait matrix. (D) Pairwise bait Pearson correlations. (E) Protein cliques (fully connected subgraphs of ≥3 baits).

**Supplemental Table 2.** AP-RNA-seq data. (A) Co-purification FPKM values (WT baits). (B) Max-normalized enrichment. (C) Hierarchically clustered, bait-averaged enrichment. (D) Per-gene nuclear processing dwell scores. (E–I) Non-coding RNA enrichment by class: CUTs, lncRNAs, SUTs, XUTs and UTR-derived transcripts.

**Supplemental Table 3.** Strains, oligonucleotides and reagents. (A) Yeast strains. (B) PrA-tagging primers. (C) Plasmids. (D) Cloning and qRT-PCR oligonucleotides. (E) Northern blot probes. (F) Affinity-purification buffers. (G) Single-molecule FISH (smFISH) probes.

**Supplemental Table 4.** Total RNA-seq of sequential mCherry-4xMS2 isolations with Cbp80p-PrA and Nab2p-PrA: per-gene coverage and FPKM (related to Figure 4F).

**Supplemental Table 5.** Label-free MS quantification of MS2CP-GFP/RNP isolations. (A–Q) Label-free quantification per isolation; the Index sheet lists each sample’s bait, promoter, transcript and the figure it supports.

**Supplemental Table 6.** Protein composition and copy number of the mCherry-4xMS2 mRNP by label-free MS (related to Figure 5A and Supplemental Figure 10F).

## DECLARATION OF GENERATIVE AI AND AI-ASSISTED TECHNOLOGIES IN THE WRITING PROCESS

During the preparation of this work the authors used Claude (Anthropic) to check analyses against the data and scripts and for text editing. After using this tool, the authors reviewed and edited the content and take full responsibility for the content of the publication.

## ACKNOWLEDGMENTS

We thank the proteomics facilities of the Institute for Systems Biology and The Rockefeller University, Dr. Denis Faubert (IRCM proteomics platform), and N. Buchler, F. Cross and R. Singer for plasmids.

This work was supported by the Canadian Institutes of Health Research (PJT-153313 to M.O.; PJT-192036 to D.Z.; PJT-174987 to M.L.A.), by the Natural Sciences and Engineering Research Council of Canada (to M.L.A, and RGPIN-2020-06924 and RGPIN-2020-00019 to M.O.), and by the National Institutes of Health (P50 GM076547 to J.D.A.; U54 GM103511, R01 GM112108 and P41 GM109824 to J.D.A. and M.P.R.). I.A. was supported by an Ontario Graduate Scholarship, a Queen Elizabeth II Graduate Scholarship in Science and Technology and scholarships from NSERC.

## AUTHOR CONTRIBUTIONS

M.P.R., P.C.F. and M.O. conceived the study. P.C.F., K.E.W., L.C.A., P.R. and M.O. performed experiments; K.E.W., E.Y.J., R.S.R., R.A.S. and R.L.M. performed mass spectrometry; I.A. and A.V.R. performed computational analyses; D.Z. and P.R. designed and analyzed smFISH. M.P.R., J.D.A., B.T.C., D.Z., M.L.A. and M.O. supervised the work, and J.D.A., P.C.F., M.P.R. and M.O. wrote the paper with input from all authors.

## METHODS

### Survey data methods Yeast strains and tagging

*S. cerevisiae* strains used in this study are listed in Supplemental Table 3A. All survey bait proteins were tagged C-terminally at their endogenous genomic loci in the W303 background (*MATa*, *ade2-1*, *ura3-1*, *his3-11,15*, *trp1-1*, *leu2-3,112*, *can1-100*) by one-step PCR-based integration of a Protein A (PrA) tag using the pMO-PrA KanMX6 plasmid as described in ^111^. Tagging cassettes were amplified using primers carrying 5’ overhangs complementary to the genomic sequences flanking each stop codon (forward overhang: 5’-CGTACGCTGCAGGTCGAC-3’; reverse overhang: 5’-ATCGATGAATTCGAGCTCG-3’) fused to gene-specific sequences. Primers, strains, and plasmids used in this study are listed in Supplemental Table 3A–C. Correct chromosomal integration was verified by colony PCR and immunoblot with horseradish peroxidase-conjugated rabbit IgG (Sigma; 1:5,000 in 5% skim milk in TBS-Tween) as described in ^111^. Signal was detected by enhanced chemiluminescence (ECL).

### Antibody-coupled affinity purification for mass spectrometry (AP-MS)

AP-MS was carried out as described in ^35,111^. Yeast cells were grown to mid-log phase (OD₆₀₀ 0.8–1.0) in YPD at 30°C and harvested by centrifugation. Cell pellets were snap-frozen in liquid nitrogen and cryogenically disrupted using a freezer mill (SPEX SamplePrep). Cryogenic lysates were resuspended in a purification buffer empirically optimized per bait (buffer type—TBT or RNP—and NaCl concentration; see Supplemental Table 3F). The TBT buffer contained 20 mM HEPES-KOH pH 7.5, 110 mM KOAc, variable NaCl (25–150 mM), 0.5% Tween-20, 0.1% NP-40, 1 mM DTT, protease inhibitors (Complete EDTA-free, Roche), and RNase inhibitor (SuperRNasin, Ambion; 1:5,000). The RNP buffer contained 20 mM HEPES-KOH pH 7.5, variable NaCl (0–150 mM), 0.5 mM MgCl₂, 0.5% Tween-20, protease inhibitors (Complete EDTA-free, Roche), and RNase inhibitor (SuperRNasin, Ambion; 1:5,000). Buffer type and NaCl concentration were empirically optimized per bait (see Supplemental Table 3F). For Yra1p and Yra2p, heparin (0.5 mg/mL) was added to TBT; for Pml39p, KOAc was adjusted to 90 mM and heparin (0.5 mg/mL) was included. Cleared lysates (15,000 × g, 15 min, 4°C) were incubated with rabbit IgG-conjugated magnetic beads (1 h, 4°C, rotation). Beads were washed four times in optimized purification buffer, followed by a 5 min wash in Last Wash buffer (0.1 M NH₄OAc, 0.1 mM MgCl₂, 0.02% Tween-20). For mass spectrometry, beads were washed four additional times in Last Wash buffer lacking Tween-20 to avoid detergent carryover. Bound complexes were eluted twice with 500 µl 0.5 M NH₄OH, 0.5 mM EDTA (20 min, room temperature); eluates were pooled and dried by vacuum centrifugation.

### Mass spectrometry for mRNP survey

Dried eluates were digested in solution with trypsin (recombinant Trypsin, Promega; overnight, 37°C) and analyzed by nanoLC-MS/MS on a Thermo LTQ Orbitrap at the Institute for Systems Biology (Seattle) using data-dependent acquisition. Each bait was purified in a minimum of two independent biological replicates, and untagged strains were processed in parallel as negative controls. Cells were grown in YPD; the analysis reported here is label-free and based on adjusted spectral counts (see AP-MS data analysis). Spectra were converted to mzXML and searched with X! Tandem (k-score) against a *Saccharomyces cerevisiae* ORF database (yeast_orfs_all, 20060126), with trypsin specificity, up to one internal missed cleavage and fully tryptic termini; carbamidomethylation (C) was set as a fixed modification, and oxidation (M) as a variable modification. Peptide-and protein-level identifications were assigned using PeptideProphet and ProteinProphet within the Trans-Proteomic Pipeline.

### AP-MS data analysis for mRNP survey

Raw MS data were processed using the ABACUS pipeline^112^ to yield adjusted spectral counts (adjSC) per prey protein per bait purification. Prey proteins were retained if adjSC > 0 and SAINT score ≥0.5^113^. Prey subcellular localization was obtained from the Saccharomyces Genome Database (SGD)^114^; only nuclear-localized prey proteins were retained. For baits that also function in the cytoplasm (e.g., Dbp5p, Gfd1p, She2p), this captures only the nuclear portion of their interactome. The resulting nuclear-filtered matrix (380 nuclear-annotated preys × 34 baits) was used for the AP-MS network analyses (Figures 1C, D, F and G, Supplemental Figures 1–3). The multi-omics integration (Figures 1E and 3A; Supplemental Figure 6) used the averaged AP-MS and AP-RNA-seq bait–bait correlations across the 18 baits profiled by both AP-MS and AP-RNA-seq (see Multi-omics hierarchical clustering). Bait-bait Pearson correlations were computed across the spectral-count profiles of all nuclear preys; the multi-omics module network (Supplemental Figure 6D) used a combined correlation threshold of r > 0.35 to define edges.

### Network topology analysis

Bait–prey and bait–bait networks were built from the nuclear-filtered AP-MS matrix using networkx (Python). Node degree per bait (Supplemental Figure 1C) was defined as the number of distinct nuclear prey proteins detected (spectral count > 0) in that bait’s purification. Protein cliques (Figure 1G; Supplemental Table 1E)—fully connected subgraphs of three or more baits that mutually co-purify one another—were enumerated from the bait co-detection graph, retaining non-nested cliques of size ≥3. For the directed network (Supplemental Figure 3E), an edge A→B was drawn when protein B was detected as a prey in the purification of bait A, weighted by the corresponding spectral count; betweenness centrality (Supplemental Figure 3F) was computed on the unweighted directed graph (networkx, normalized). For the undirected co-purification network (Supplemental Figure 3G), two baits were connected when either detected the other, and node degree was used to identify the most broadly connected hub proteins. Pairwise weighted shortest-path distances between baits (Supplemental Figure 3A–D) were computed by Dijkstra’s algorithm on the bait–prey bipartite graph, with edge cost defined as 1/log₁₀(spectral count + 1) so that stronger co-purification corresponds to a shorter distance; the resulting distance matrices were ordered by pathway position and by average-linkage hierarchical clustering, and mean distances were computed between functional groups.

### Antibody-coupled affinity purification for RNA sequencing (AP-RNA-seq) for mRNP survey

AP-RNA-seq was performed as described in^2^. Cryogenic lysates were prepared as for AP-MS (above). Co-purified RNA was recovered from washed bead eluates by Trizol/chloroform extraction and ethanol precipitation, ribosomal RNA was depleted, and sequencing libraries were prepared and sequenced on a HiSeq2500. Per-gene read counts were normalized to FPKM. Genes with FPKM < 200 in all bait purifications were excluded. For each gene, enrichment per bait was normalized to the per-gene maximum across all baits (max-FPKM normalization; range 0–1). Eighteen wild-type bait purifications encompassing 2,452 mRNA genes were used. AP-RNA-seq data were also compared to the published Mlp1p AP-RNA-seq dataset^2^. Mlp1p association was taken as the mean normalized read count of the basket-containing Mlp1p purification minus that of its IgG control, and preference for basket-containing over basket-less NPCs as the DESeq2 log2 fold change between the two purifications. Spearman correlations of each bait’s enrichment with both measures were computed across the 2,431 genes shared with the 18-bait survey (2,419 with a fold change), and the fold change of each transcript cluster was compared with that of all other clusters by two-sided Mann–Whitney U test.

### AP-RNA-seq data analysis for mRNP survey

Per-bait mean enrichment was computed across replicates. Hierarchical clustering of the co-enriched coding genes (max-normalized enrichment > 0.3 in at least one bait; n = 2,434 of the 2,452-ORF set; Figure 2A) used Ward linkage on Pearson correlation distance (1 − r), and the dendrogram was cut into k = 8 clusters; for display in Figure 2A, genes were ordered by average-linkage clustering on the same distance. A nuclear processing dwell score was computed for each gene as the sum, over the eight pathway stages, of its highest mean enrichment among the baits assigned to each stage (dwell AUC; maximum 8; stages and their baits: CBC, Cbp80p; THO/TREX, Sub2p and Thp2p; splicing, Snu71p; hnRNP, Nab2p, Hrp1p, Npl3p, Pab1p and Gbp2p; 3′-end processing, Clp1p, Syc1p and Ref2p; TREX-2/NPC, Sac3p and Gfd1p; export, Mex67p, Dbp5p and Yra1p; surveillance, Rrp6p), a proxy for the breadth and persistence of co-purification with the biogenesis machinery that correlates closely with the count of stages enriched above 0.3 (Spearman ρ = 0.92). The full max-normalized enrichment matrix, which additionally includes non-coding RNA classes (CUTs, SUTs, XUTs, lncRNAs and UTR-derived transcripts), comprises 9,136 transcripts (Supplemental Table 2); the Figure 2 clustering was restricted to coding ORFs. Transcript length quartiles (Q1 < 598 nt, Q2 598–960 nt, Q3 960–1,529 nt, Q4 > 1,529 nt) were defined on the 2,180 genes of the 2,452-ORF set with annotated transcript length; Spearman correlations between length and per-bait enrichment were computed (all 18 baits significant at p < 0.001). Gene Ontology enrichment of each transcript cluster, and of the transcripts co-enriched by each bait (mean normalized score > 0.3), was tested against all 2,447 AP-RNA-seq coding ORFs carrying a biological-process GO-slim annotation (SGD GO-slim mapping) by one-sided Fisher’s exact test, with Benjamini–Hochberg correction across all tests. Nab2p and Pab1p enrichment was compared within each transcript cluster by two-sided paired Wilcoxon signed-rank test with Benjamini–Hochberg correction across the eight clusters, and the overlap of their co-enriched transcript sets (mean normalized score > 0.3) by hypergeometric test. The Nab2p values derive from the survey purification; the cluster differences reported were confirmed in an independent series of three Nab2p and three Pab1p purifications. Promoter class analysis used Rhee and Pugh (2012)^115^ Supplementary Data 1 to classify 1,878 of 2,452 genes (77%) as TATA-containing/TATA-less and TAF1-enriched/TAF1-depleted; pairwise dwell-score comparisons used Mann–Whitney U tests with Bonferroni correction (six comparisons; α = 0.05/6 = 0.0083). Cross-dataset concordance was assessed by comparing bait-bait AP-MS Pearson correlations with bait-bait AP-RNA-seq Spearman correlations (thresholds: AP-MS Pearson r > 0.35 and AP-RNA-seq Spearman ρ > 0.50, met by 21 and 45 of the 153 bait pairs, respectively). A rank correlation was used for AP-RNA-seq because max-normalized FPKM values are bounded and skewed. For RNA-class comparisons (Figure 2F), coding ORFs, snRNAs and snoRNAs were assigned using the SGD annotation (R64), and CUTs, SUTs and XUTs using the annotations applied for quantification; for each bait, the median replicate-mean FPKM of each class relative to that of coding mRNAs was compared with the same ratio in total RNA from wild-type (W303) cells (six libraries) and is shown as log₂ enrichment. lncRNA-annotated features (2014 annotation) were not analyzed as a separate class because, with every bait, they co-purified in the same proportion to mRNA as in total RNA.

### Multi-omics hierarchical clustering

Throughout, ‘cluster’ denotes a data-driven grouping (hierarchical clustering or bootstrap analysis; e.g., the transcript clusters C1–C8), whereas ‘functional group’ (a fine pathway category) and ‘functional module’ (one of the four coarse multi-omics groupings) denote biologically assigned groupings; functional modules are assigned by pathway role and are displayed against the data-driven clustering, which they need not match. AP-MS and AP-RNA-seq data were integrated across the 18 shared baits by computing a bait–bait Pearson correlation independently for each readout—over the 380 nuclear preys (AP-MS) and 2,434 co-enriched transcripts (AP-RNA-seq), then averaging the two correlation matrices element-wise. Hierarchical clustering used 1−Pearson’s r distance and average linkage; robustness was assessed by multiscale bootstrap (approximately unbiased [au] p-values^116^), resampling the features of both readouts across ten scales (1,200 resamples per scale) and re-averaging the correlation. Support for the principal branches is shown in Supplemental Figure 6A. Within- and between-module correlations of the four functional modules were compared over all bait pairs by one-sided Mann–Whitney U test (Figure 1E).

### Chromosome interaction domain (CID) analysis

CID boundary coordinates and boundary-strength scores were taken from Hsieh et al. (2015)^55^, their Supplemental Table S4 (Micro-C consensus CID boundary calls; strength range 1.01–13.85, median 4.19). Gene coordinates were parsed from the SGD GFF3 annotation (R64), retaining features of type gene with systematic ORF identifiers; chromosome labels were converted to integers to match the boundary table and gene midpoints computed as (start + end)/2. Of 2,452 AP-RNA-seq genes, 2,445 (99.7%) were matched to GFF3 coordinates. For each gene, flanking boundaries on the same chromosome were located with numpy.searchsorted. The gene was assigned to the enclosing inter-boundary interval (CID), and distance to the nearest boundary computed as min(midpoint − left boundary, right boundary − midpoint); genes in intervals > 50 kb (chromosome ends lacking flanking boundaries) were excluded, retaining 2,443 genes (median CID size 4.1 kb). Associations of boundary distance with RNA cluster, nuclear dwell-score quartile and promoter class were tested by Kruskal–Wallis H (omnibus) followed by pairwise Mann–Whitney U with Bonferroni correction, and by Spearman correlation for continuous variables. Boundary strength (Hsieh 2015 score)^55^ was correlated (Pearson) with the mean dwell score of genes within 5 kb of each boundary (Supplemental Figure 8C), and the fraction of SAGA-regulated genes within 5 kb of each boundary was correlated (Pearson) with their mean dwell score (Supplemental Figure 8D). Correlations were reported individually without correction.

### Statistical analyses for mRNP survey

Non-parametric tests were used throughout because distributional normality was not assumed: Mann–Whitney U and Kruskal–Wallis tests for group comparisons (scipy.stats); Bonferroni correction for multiple comparisons unless noted; Spearman or Pearson correlations, as stated for each analysis. Analyses were performed in Python 3.11 (numpy 1.24, pandas 1.5, scipy 1.10, matplotlib 3.7, networkx 3.1, scikit-learn 1.3). Hierarchical clustering and multiscale-bootstrap validation were performed in Python (scipy 1.10.x). Analysis code is available at GitHub (https://github.com/oeffingerlab/mRNP-survey-yeast).

### Cloning

The *MET3* promoter was subcloned from p404MET3 into pRS414 using SacI and SpeI restriction sites to create pRS414-MET3. HA-MS2(ΔFG)-eGFP-SV40NLS was PCR-amplified from pMS2-GFP^117^ and cloned into pRS414-MET3 to create pRS414-MET3-MS2CP-GFP. Wild-type eGFP and mCherry sequences were PCR-amplified and cloned into pRS426-TDH3 or pRS426-GAL1 using SpeI and XmaI restriction sites. Four repeats of the MS2 hairpin were inserted after these sequences between XmaI and XhoI sites, using complementary primers (Supplemental Table 3D). Equivalent primers with SpeI restriction sites were used to insert the 4xMS2 upstream of mCherry for 5’ tagging. Primers with 1 or 2 MS2 hairpin repeats were used for 1x or 2x versions. 6xMS2 was subcloned from pSL-MS2-6x and 24xMS2 from pCR4-24XMS2SL-stable^88^.

*ACT1* and *RPS30B* intron sequences were amplified from yeast W303 genomic DNA (primers in Supplemental Table 3D), and then blunt-end ligated into pRS426-mCherry-4xMS2 and pRS426-mCherry plasmids digested with MscI (31 bp into the mCherry CDS). The *INO1* promoter was PCR amplified from yeast genomic DNA and subcloned into the pRS426-mCherry and pRS426-mCherry-4xMS2 plasmids using SacI and SpeI restriction sites (primers in Supplemental Table 3D). The *ASH1* 3’ UTR sequence was amplified from yeast genomic DNA and subcloned into pRS426-mCherry and pRS426-mCherry-4xMS2 plasmids after the 4xMS2 tag using XhoI and EagI restriction sites (primers in Supplemental Table 3D).

### Strains for sequential RNA affinity purifications

Strains used in this work were derived from *S. cerevisiae* wild-type W303 (*MATa ade2-1 ura3-1 his3-11,15 trp1-1 leu2-3,112 can1-100*). *NAB2*, *CBP80*, *CDC33*, *MEX67*, and *THP2* were genomically tagged at the C-terminus using a Protein A::*HIS5* cassette with a 3C protease site (GLEVLFQGPS) inserted at the 5’ end^118^. For sequential RNA purifications, pRS414-MS2CP-GFP was transformed into the appropriate strain, followed by transformation with the transcript-encoding pRS426 plasmid. Transformations were performed using standard lithium acetate techniques. Cells were grown at 30°C in complete synthetic defined medium lacking L-tryptophan, uracil, and L-methionine.

### Sequential RNA affinity purifications

Yeast cells were grown, harvested at mid-log phase and cryogenically milled as previously described, and affinity isolations performed by adapting previously reported methods^111,119^. 2 g of frozen cell powder was resuspended in 8 ml of RNP buffer (20 mM HEPES, pH 7.4; 150 mM NaCl; 110 mM K-acetate; 1 mM DTT; 0.1% Tween 20; 0.5% Triton X-100; 1:5000 RNasin (Promega); 1:5000 Antifoam B; 1 mM PMSF; 3 µg/ml pepstatin A). Where indicated, complexes were stabilized by a short in-lysate glutaraldehyde fixation before capture: glutaraldehyde was added to the resuspended grindate to a final concentration of 3 mM (0–3 mM for the titration in Supplemental Figure 10E) and incubated for 5 min. on ice, before the reaction was quenched with Tris-HCl pH 8 to a final concentration of 100 mM. A parallel isolation without glutaraldehyde was carried out from the same material for comparison. Glutaraldehyde concentrations were optimized for the complexes to be stabilized; the procedure stabilizes dynamic associations during purification and establishes how much of the composition is lost without it, rather than mapping protein–protein contacts^2,70,120^. Lysate was spun at 10,000 x g for 10 min., and the supernatant incubated with a 200 µl suspension (30 mg beads) of IgG-conjugated Dynabeads equilibrated in RNP buffer (Thermo Fisher). After a 30 min. incubation at 4°C, beads were washed 3 times with 1 ml RNP buffer and resuspended in 150 µl RNP with 80 ng/µl PreScission Plus (HRV-3C) protease. This was incubated for 1 hr at 4°C, and the supernatant was collected. This was pooled with an additional 75 µl RNP wash of the beads, and the combined solution was incubated for 30 min with a 10 µl suspension (1.5 mg beads) of Dynabeads conjugated to LaG16-G4S-LaG2 nanobody^62^. After 3 washes with RNP buffer, protein was eluted at 72°C for 10 min. with 25 µl NuPAGE LDS buffer. When prepared for MS analysis, this elution was reduced for 10 min. at 72°C with 25 mM DTT, then alkylated with 0.1 M iodoacetamide for 30 min. at room temperature. For RNA isolation, beads were instead resuspended in 600 µl RLT buffer for RNeasy purification (Qiagen).

To determine yields, samples of resuspended lysate were taken before and after each binding step. These were run on a 4-12% Novex Bis-Tris gel in MES running buffer (Thermo Fisher), and probed by Western blotting using, for GFP, mouse anti-GFP antibody (Roche, cat. no. 11 814 460 001) diluted 1:1,000 in TBST / 2% dry milk and an anti-mouse, HRP-conjugated secondary (GE Healthcare, cat. no. NA931V) diluted 1:3,000 in TBST / 2% dry milk. For PrA, membranes were instead probed with peroxidase anti-peroxidase (Sigma, cat. No. P1291) diluted 1:5,000 in TBST. Signals were quantified using ImageJ software.

### Mass spectrometry-based proteomic analysis of mCherry-MS2-associated proteins

Alkylated affinity purification elutions were purified as single gel plugs by SDS-PAGE: samples were run at 125 V for 7 min. on a 10% Bis-Tris gel (Thermo Fisher), which was then stained with Coomassie Blue dye. Bands were then excised as a single piece, destained, and dehydrated. In-gel digestion with trypsin was performed on destained dehydrated gel pieces overnight with 0.5-1 µg trypsin in 50 µl 50 mM ammonium bicarbonate (ABC) buffer with 10% acetonitrile (ACN). Tryptic peptides were extracted with POROS20 C18 beads^121^, desalted on Zip Tips (0.6 µl C18 resin, Millipore), sequentially eluted with 40% and then 90% acetonitrile / 0.1% acetic acid, and the eluates pooled, concentrated by vacuum centrifugation, and suspended in 10 µl of 0.1% formic acid (FA). Desalted peptides (2 µl) were analyzed by direct injection onto a C18 analytical column (EASY-Spray 3 µm, 150 mm x 0.075 mm) using a Thermo EASY-nLC 1000 nanoLC coupled online to an Orbitrap Q Exactive Plus (QE+) or Fusion mass spectrometer (Thermo Fisher). Peptide separation and MS detection was performed over 60 min at ∼300 nl/min using a discontinuous ACN gradient from 0 to 30 % mobile phase B over 60 min, followed by 30 to 100% B over 15 min (mobile phase A, 0.1% FA; mobile phase B, 97% ACN in 0.1% FA). The spray voltage was set at 1.9–2.3 kV. The instruments were operated in the data-dependent mode, in which the most abundant ions were fragmented by higher-energy collisional dissociation (HCD) (HCD collision energy 30 (QE+) or 35 (Fusion)) and analyzed in the Orbitrap mass analyzer. The target resolution for MS1 was 70,000 (QE+) and 120,000 (Fusion), and for MS2 was 17,500 (QE+) and 30,000 (Fusion). Ions (300–1,700 m/z QE+, 400-1500 m/z Fusion) with a charge state of >1 were selected for fragmentation. A dynamic exclusion of 10 s (QE+) and 30 s (Fusion) and a ‘lock mass’ at 371.1012 Da were used.

### Mass spectrometry data processing and quantitation for Sequential RNA affinity purifications

Combined MS/MS spectra were extracted from Thermo RAW files derived from three biological replicate samples and searched by Proteome Discoverer/SEQUEST (version 2.0/2.1, Thermo Fisher) against a *Saccharomyces cerevisiae* protein sequence database and a common contaminants database. SEQUEST search parameters were as follows: full trypsin specificity with 2 missed cleavages, precursor and fragment tolerances of 20 ppm and 0.6 Da, fixed modification of carbamidomethylation of cysteine, and variable modification for methionine oxidation. Putative SEQUEST peptide-spectrum matches (PSMs) were scored by the Percolator algorithm^122^ to assign individual expectation scores (PEP). PSMs were filtered to achieve an estimated 1% peptide false discovery rate (q-value < 0.01) and then PSMs were assembled into protein groups using the “strict maximum parsimony principle” option. Relative label-free quantitation (LFQ) was performed using the Proteome Discoverer precursor ions area quantification workflow on the top 3 most intense peptides per protein. Results were exported to Excel for downstream analysis. Abundances for the three biological replicates were averaged. Proteins with fewer than three peptides were not analyzed. For per-transcript copy-number estimates (Supplemental Table 6), LFQ protein abundances were normalized to MS2CP-GFP, assuming two MS2CP-GFP copies per MS2 hairpin (one coat-protein dimer per hairpin). For other relative comparisons, abundances were normalized to the corresponding PrA-tagged bait protein.

### RNA-seq and real-time PCR for Sequential RNA affinity purifications

RNA from sequential affinity purifications was purified using the RNeasy kit (Qiagen). RNA was fragmented and ligated to adaptors before library preparation using Illumina kits and run on a HiSeq 2500 sequencer with 50 bp single reads. Reads were mapped to *S. cerevisiae* exons or mCherry-4xMS2 using BBMap v39.08^123^ and quantified with StringTie v3.03^124^.

For qRT-PCR, RNA was reverse transcribed using SuperScript VILO (Thermo Fisher). Real-time PCR was then performed using PowerUp SYBR Green mix (Thermo Fisher) on a QuantStudio 12K Flex system using standard conditions (Thermo Fisher). Primer pairs used to quantify each gene are listed in Supplemental Table 3D^125^.

### Northern hybridization

RNA samples were separated on a 6% TBE-Urea acrylamide gel, and Northern blotting was performed as previously described^126^. Briefly, gels were transferred to Hybond N+ membranes (GE Healthcare) and hybridized at 37°C overnight with DNA oligomer probes (listed in Supplemental Table 3E) that were end-labeled with γ^32P^-ATP and diluted in a buffer of 50% formamide, 5x SSPE, 5x Denhardt’s buffer, 0.2 mg/ml fish sperm DNA, and 0.2% SDS. Membranes were then washed twice for 15 min. at 42°C with 6xSSPE before being exposed to a Phosphorimager screen.

### Sucrose density gradient centrifugation

Cells expressing MS2CP-GFP together with mCherry-4xMS2, with or without the RPS30B intron, in the Cbp80p-PrA background were processed as in the sequential RNA affinity isolation protocol up to the first affinity step; the MS2 capture step was omitted. After 3C protease elution from the IgG Dynabeads, 150 µl of each digest was loaded on top of a 10-40% sucrose gradient in a buffer of 20 mM HEPES, pH 7.4, 150 mM NaCl, 110 mM K-acetate, 1 mM DTT, 0.1% Tween 20, 0.5% Triton X-100, 1:5000 RNasin (Promega), 1x Protease Inhibitor Cocktail (Sigma). Gradients were centrifuged using an SW 55 TI rotor (Beckman Coulter) at 40,000 rpm for 4.5 hrs at 5°C. Gradients were manually unloaded from the top in 23 fractions of 205 µl. 12 µl samples were then separated on 4-12% Bis-Tris gels, run in duplicate for SYPRO Ruby staining or anti-GFP Western blotting. The SYPRO signal reports the total Cbp80p-associated population, whereas the anti-GFP signal follows MS2CP-GFP and thereby the MS2-tagged reporter mRNP; the sedimentation profiles in Figure 6D were quantified from the anti-GFP blots. Sedimentation coefficients were estimated from the position of each peak in the gradient using the tables of McEwen for linear sucrose gradients^127^; they were calculated rather than calibrated against sedimentation standards.

### Single-molecule fluorescence in situ hybridization (smFISH)

smFISH probe design, labeling, and hybridization to visualize mCherry reporter mRNAs were performed as described^128^. In brief, cells were grown to mid-log phase, fixed with paraformaldehyde, and digested with lyticase before immobilization on poly-L-lysine–treated coverslips. Cells were then hybridized with probes complementary to the mCherry mRNA (Supplemental Table 3G), and three-dimensional (3D) datasets were acquired on an upright wide-field microscope. For visualization, 3D image stacks were converted to 2D by maximum-intensity projection in ImageJ.

## SUPPLEMENTAL FIGURES

**Supplemental Figure 1.**
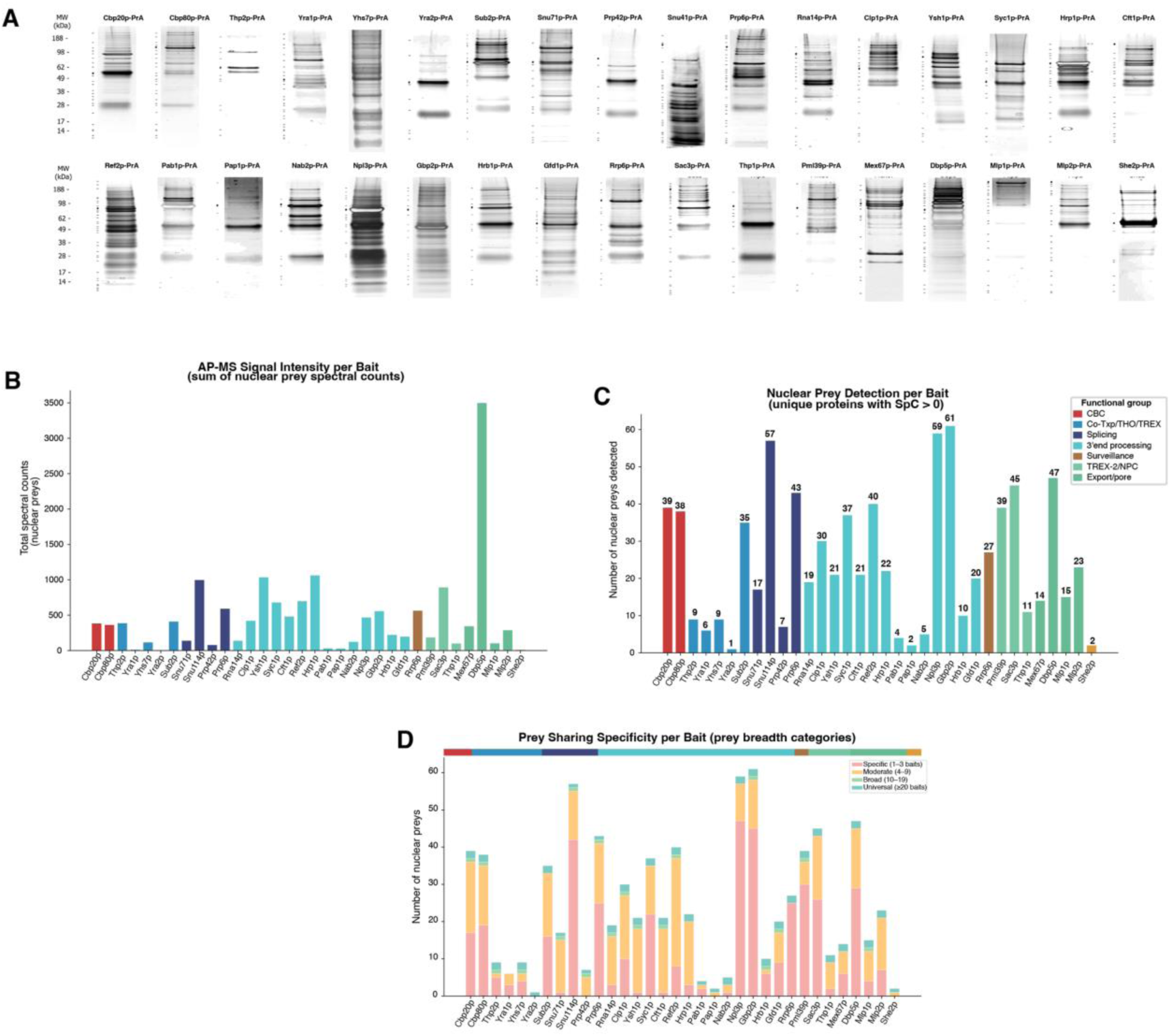
AP-MS quality controls and bait validation. **(A)** Coomassie-stained SDS-PAGE of single-step IgG purifications of each of the 34 PrA-tagged baits. **(B)** AP-MS signal intensity per bait (sum of nuclear prey spectral counts), pathway order; bars colored by functional group. **(C)** Nuclear prey detection breadth per bait (unique nuclear preys, SpC > 0). **(D)** Prey sharing per bait: number of nuclear preys (SpC ≥ 2) per bait that are specific (detected in 1–3 baits), moderate (4–9), broad (10–19) or universal (≥20); pathway order.

**Supplemental Figure 2.**
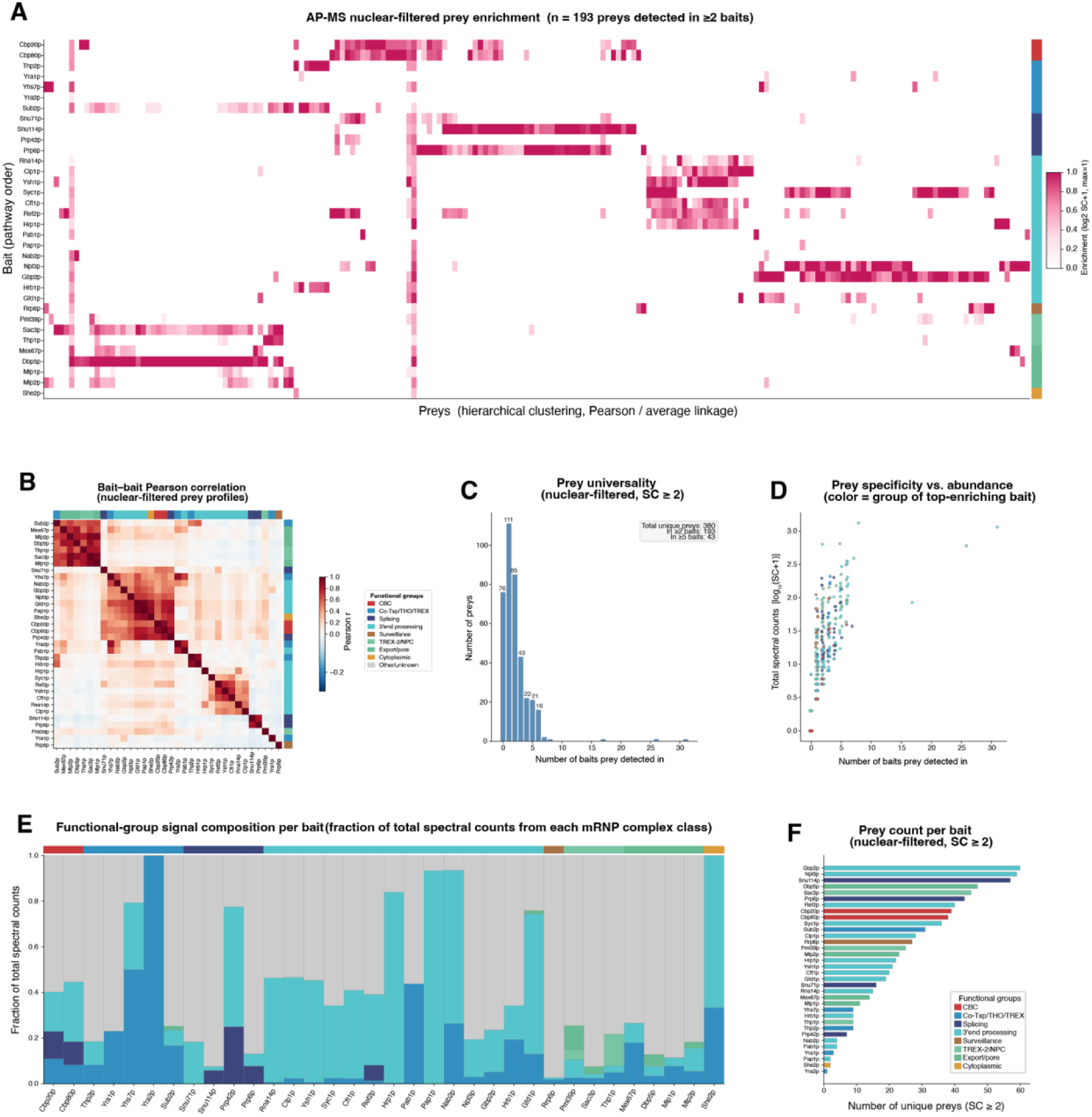
AP-MS dataset characterization. **(A)** Nuclear-filtered prey enrichment heatmap [log₂(SpC+1), per-prey normalized] for all nuclear preys in ≥2 baits; baits in pathway order (rows), preys by hierarchical clustering (columns). **(B)** Bait–bait Pearson correlation matrix, baits ordered by clustering. **(C)** Prey universality distribution: histogram of nuclear preys (SpC ≥ 2) by number of detecting baits. **(D)** Prey specificity versus abundance [log₁₀(SpC+1) vs. number of detecting baits], colored by functional group of the highest-enriching bait. **(E)** Functional-group signal composition per bait. **(F)** Prey count per bait (SpC ≥ 2), ranked; colors indicate functional group.

**Supplemental Figure 3.**
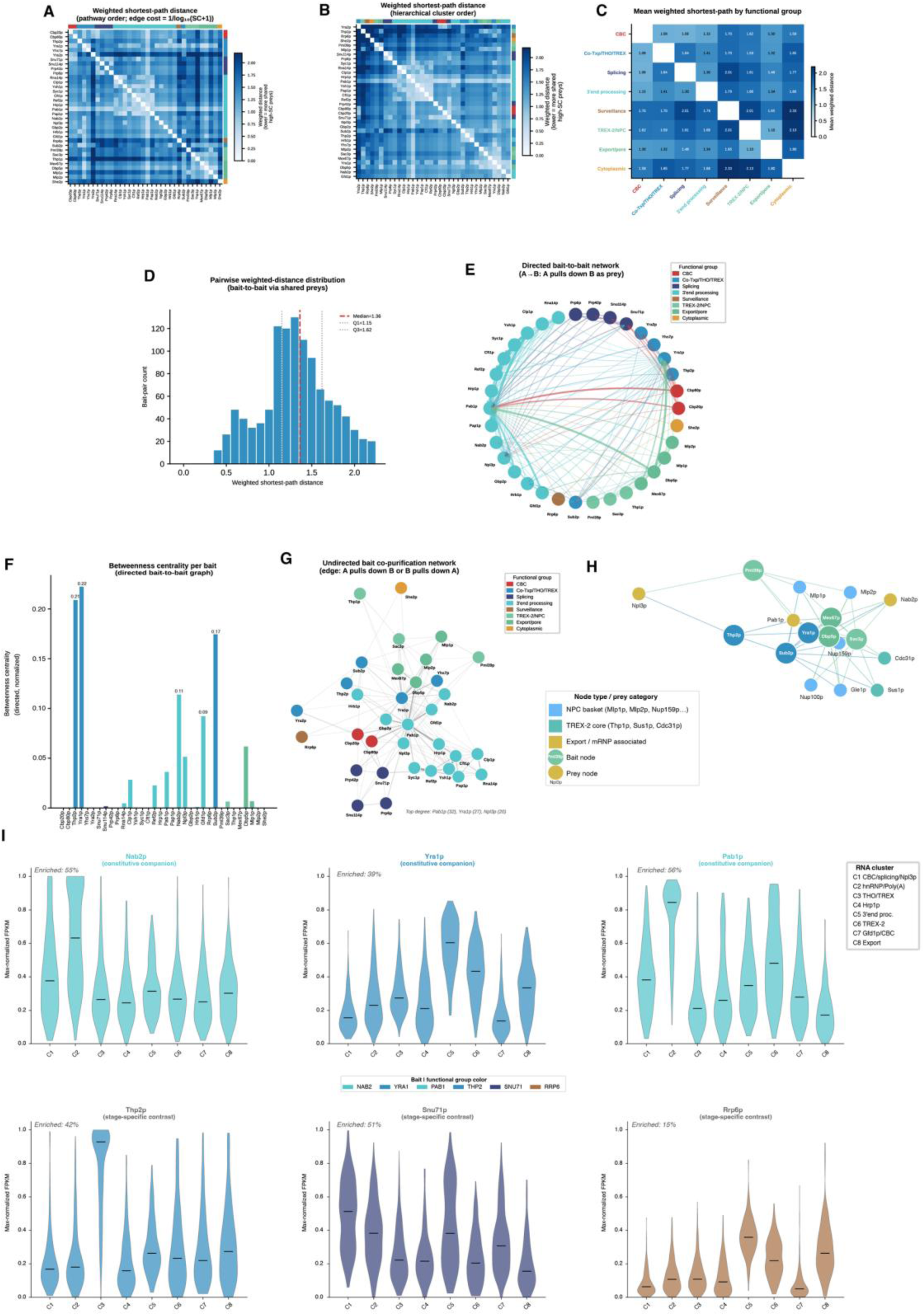
AP-MS bait interaction network topology. **(A, B)** Weighted pairwise shortest-path distances between baits (edge cost = 1/log₁₀(SpC+1)), ordered by pathway (A) and by hierarchical clustering (B); lighter = shorter path. **(C)** Mean weighted shortest-path distance between functional groups. **(D)** Distribution of pairwise weighted shortest-path distances. **(E)** Directed bait-to-bait network (edge A→B: B detected as a prey of A); nodes colored by functional group. **(F)** Betweenness centrality per bait in the directed graph. **(G)** Undirected bait co-purification network (edge: A pulls down B or B pulls down A); node color, functional group. **(H)** Pml39p-centered export scaffold: Mlp1p, Mlp2p and the NPC tether. **(I)** Transcript co-enrichment per RNA cluster for broadly associated (Nab2p, Yra1p, Pab1p) and stage-specific (Thp2p, Snu71p, Rrp6p) baits; horizontal bar, median.

**Supplemental Figure 4.**
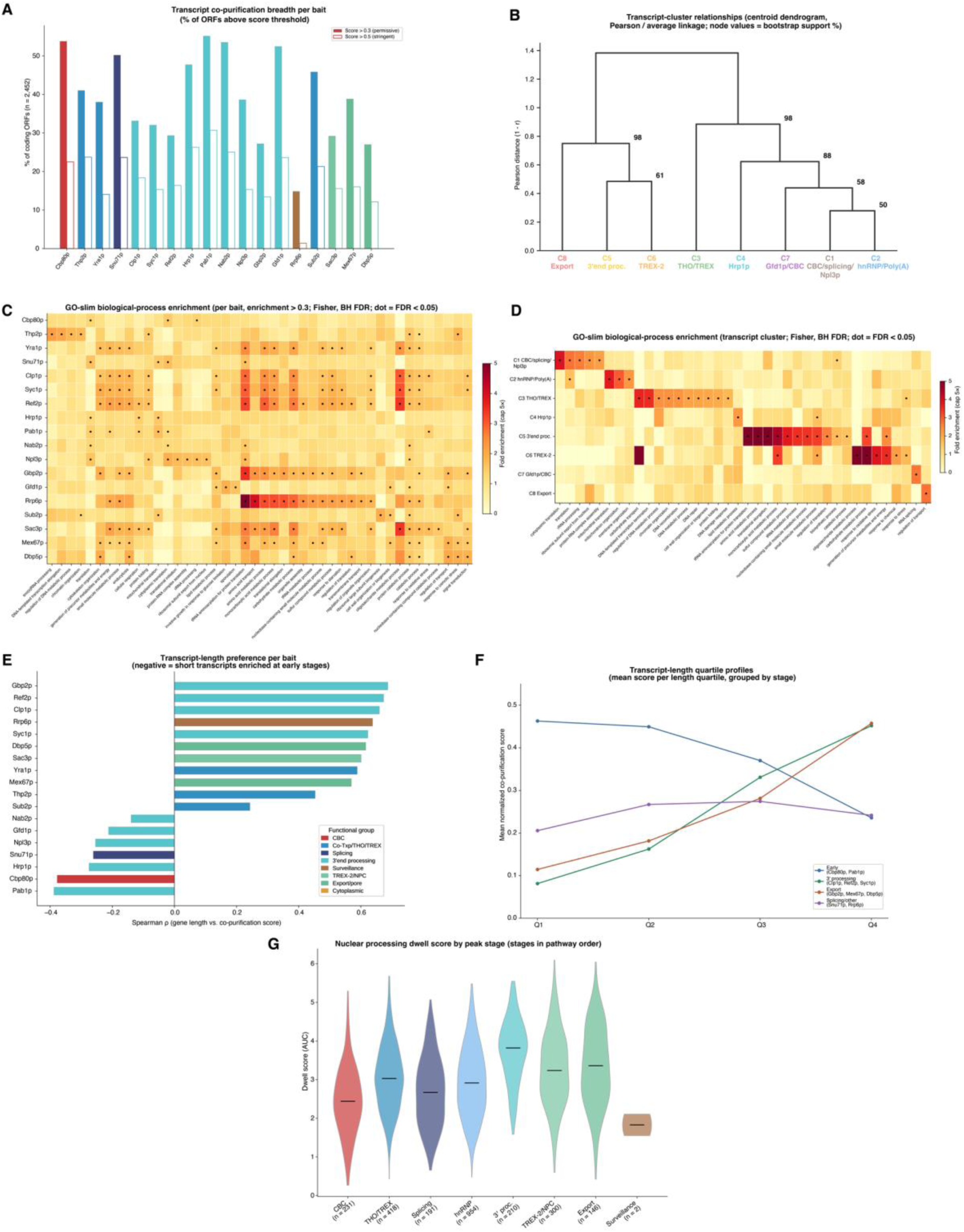
AP-RNA-seq deep characterization of transcript co-purification. **(A)** Transcript co-purification breadth per bait: percentage of covered coding ORFs (n = 2,452) with mean normalized score >0.3 (filled) or >0.5 (outline), pathway order. **(B)** Transcript-cluster relationships: centroid dendrogram of the eight transcript clusters (Pearson distance, average linkage); node values, bootstrap support (%). **(C)** GO-slim biological-process enrichment per bait: transcripts with mean normalized co-purification score > 0.3 for each bait, relative to all annotated AP-RNA-seq ORFs (n = 2,447; capped at 5×); one-sided Fisher’s exact test with Benjamini–Hochberg correction; dots, FDR < 0.05. **(D)** GO-slim biological-process enrichment per transcript cluster (C1–C8): fold enrichment relative to all annotated AP-RNA-seq ORFs (n = 2,447; capped at 5×); one-sided Fisher’s exact test with Benjamini–Hochberg correction; dots, FDR < 0.05. **(E)** Transcript-length preference per bait: Spearman ρ between transcript length and mean normalized co-purification score (n = 2,180 genes; all p < 10⁻¹⁰); negative, enrichment on shorter transcripts; positive, enrichment on longer transcripts. Scores are normalized per transcript across baits, so ρ reflects each bait’s relative length preference. **(F)** Mean normalized co-purification score per transcript-length quartile (Q1 < 598 nt, Q2 598–960 nt, Q3 960–1,529 nt, Q4 > 1,529 nt) for representative bait-stage groups. **(G)** Nuclear processing dwell score by peak biogenesis stage (the stage at which each gene is most strongly enriched), stages in pathway order (CBC, THO/TREX, splicing, hnRNP, 3′-end processing, TREX-2/NPC, export, surveillance; stage–bait assignment in Methods); number of genes per stage below each violin; horizontal bar, median.

**Supplemental Figure 5.**
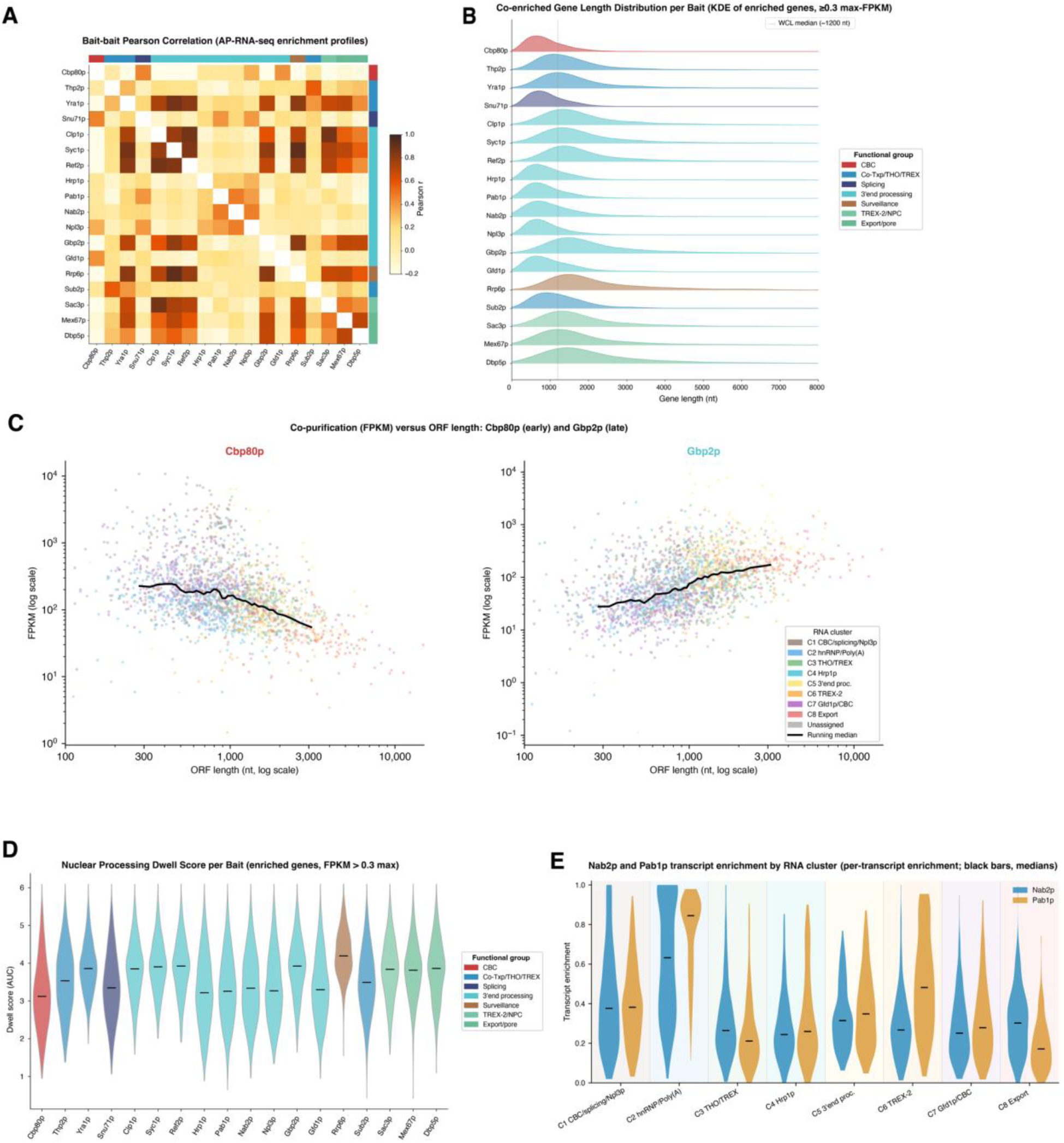
AP-RNA-seq extended analyses. **(A)** Bait–bait Pearson correlation matrix (AP-RNA-seq only). **(B)** Co-enriched gene length distribution per bait (KDE ridge plot; dashed line, 1,200 nt reference). **(C)** Co-purification enrichment (FPKM) versus ORF length for representative early-stage (Cbp80p) and later-stage (Gbp2p) baits, log–log axes; black line, running median. **(D)** Nuclear processing dwell score per bait (co-enriched genes, normalized score > 0.3), pathway order; horizontal bar, median. **(E)** Per-transcript co-purification enrichment for Nab2p (blue) and Pab1p (amber) across the eight transcript clusters (C1–C8; n = 179–505 transcripts per cluster); black bars, medians; background shading, cluster color. Paired Wilcoxon signed-rank test per cluster with Benjamini–Hochberg correction.

**Supplemental Figure 6.**
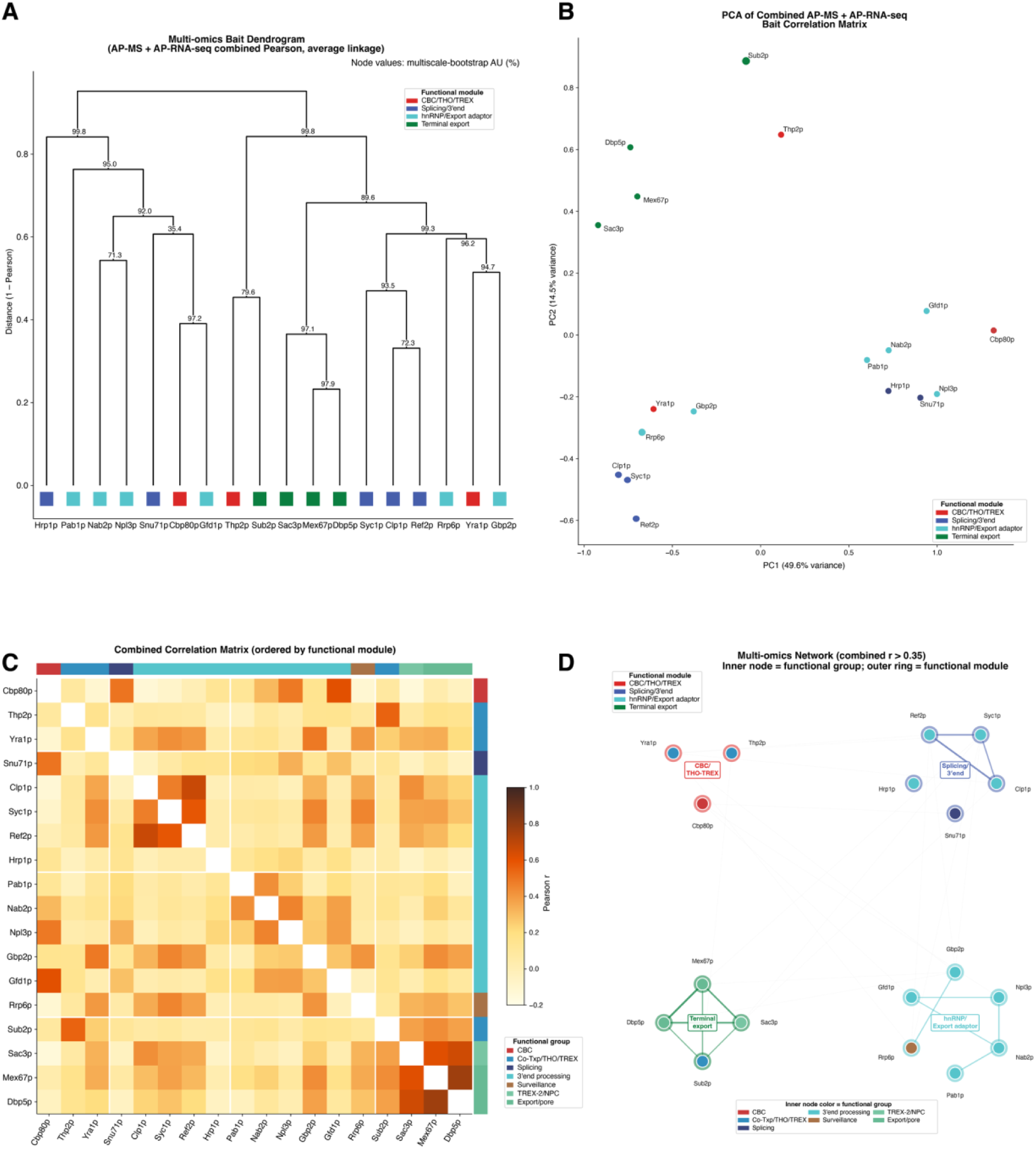
Robustness of the multi-omics functional-module assignment: four complementary views of the combined bait–bait correlation matrix (mean of the AP-MS and AP-RNA-seq Pearson correlations across the 18 shared baits; robustness by multiscale-bootstrap AU). **(A)** Bait dendrogram (combined-correlation distance, average linkage); bait names in black, with a per-bait functional-module marker beneath each leaf. Values at internal nodes are multiscale-bootstrap support (AU, %). Because functional modules are assigned by pathway role, a module’s members may fall on different branches of the data-driven tree (e.g., Sub2p groups with Thp2p rather than with the other export baits). **(B)** Principal component analysis of the same combined correlation matrix (each point a bait, colored by functional module; no enclosing polygons). **(C)** Full correlation heatmap ordered by functional module as contiguous blocks (white lines mark module boundaries); side bars indicate functional group (fine, 8 categories) and functional module (coarse, 4). **(D)** Functional-module interaction network (edges for combined r > 0.35; edge width and opacity scale with r; within-module edges colored by module, cross-module edges gray; node fill = functional group, outer ring = functional module; nodes are arranged by module for legibility).

**Supplemental Figure 7.**
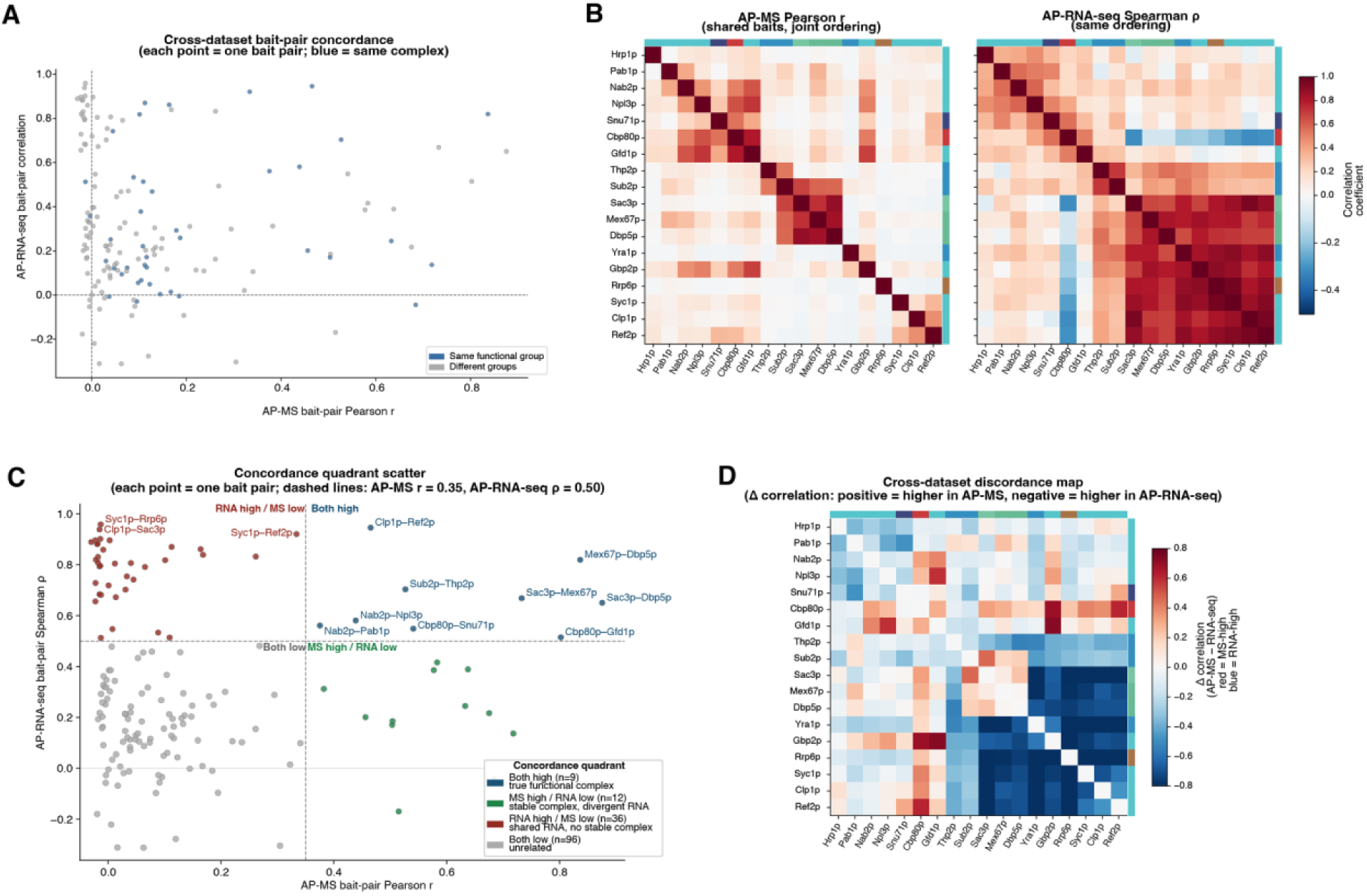
Cross-dataset concordance between AP-MS and AP-RNA-seq. **(A)** Bait-pair concordance scatter (x: AP-MS Pearson r; y: AP-RNA-seq Spearman ρ); pairs from the same functional group in blue. **(B)** Side-by-side AP-MS and AP-RNA-seq correlation heatmaps, identical clustering order; block-diagonal structure confirms shared pathway-stage architecture. **(C)** Concordance quadrants: each bait pair classified by AP-MS Pearson r > 0.35 and AP-RNA-seq Spearman ρ > 0.50 (dashed lines); both high, n = 9; MS high/RNA low, n = 12; RNA high/MS low, n = 36; both low, n = 96. **(D)** Discordance map [Δr = AP-MS Pearson r − AP-RNA-seq Spearman ρ]; red: more correlated in AP-MS; blue: more in AP-RNA-seq.

**Supplemental Figure 8.**
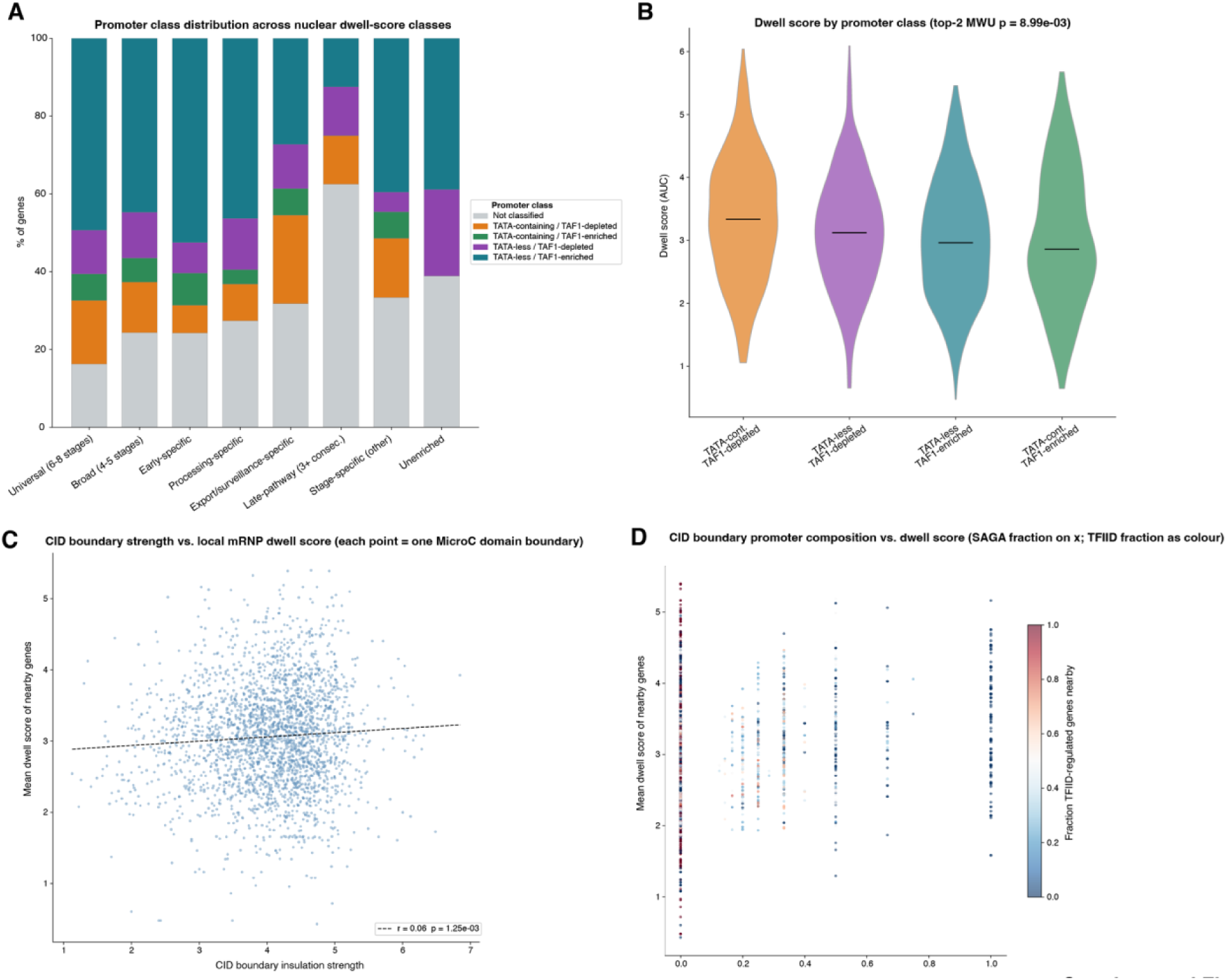
Genomic and chromatin context of mRNP co-purification. **(A)** Promoter class distribution across nuclear dwell-score classes. **(B)** Dwell score distribution by promoter class (violin plots; medians and Mann–Whitney U p-value shown). **(C)** CID boundary insulation strength versus mean dwell score of genes within 5 kb^55^; regression line and Pearson r (r = 0.06, p = 1.25×10⁻³). **(D)** CID boundary promoter composition versus mean local dwell score; color encodes TFIID-regulated fraction (Pearson r = 0.18, p = 5.9×10⁻²⁰).

**Supplemental Figure 9.**
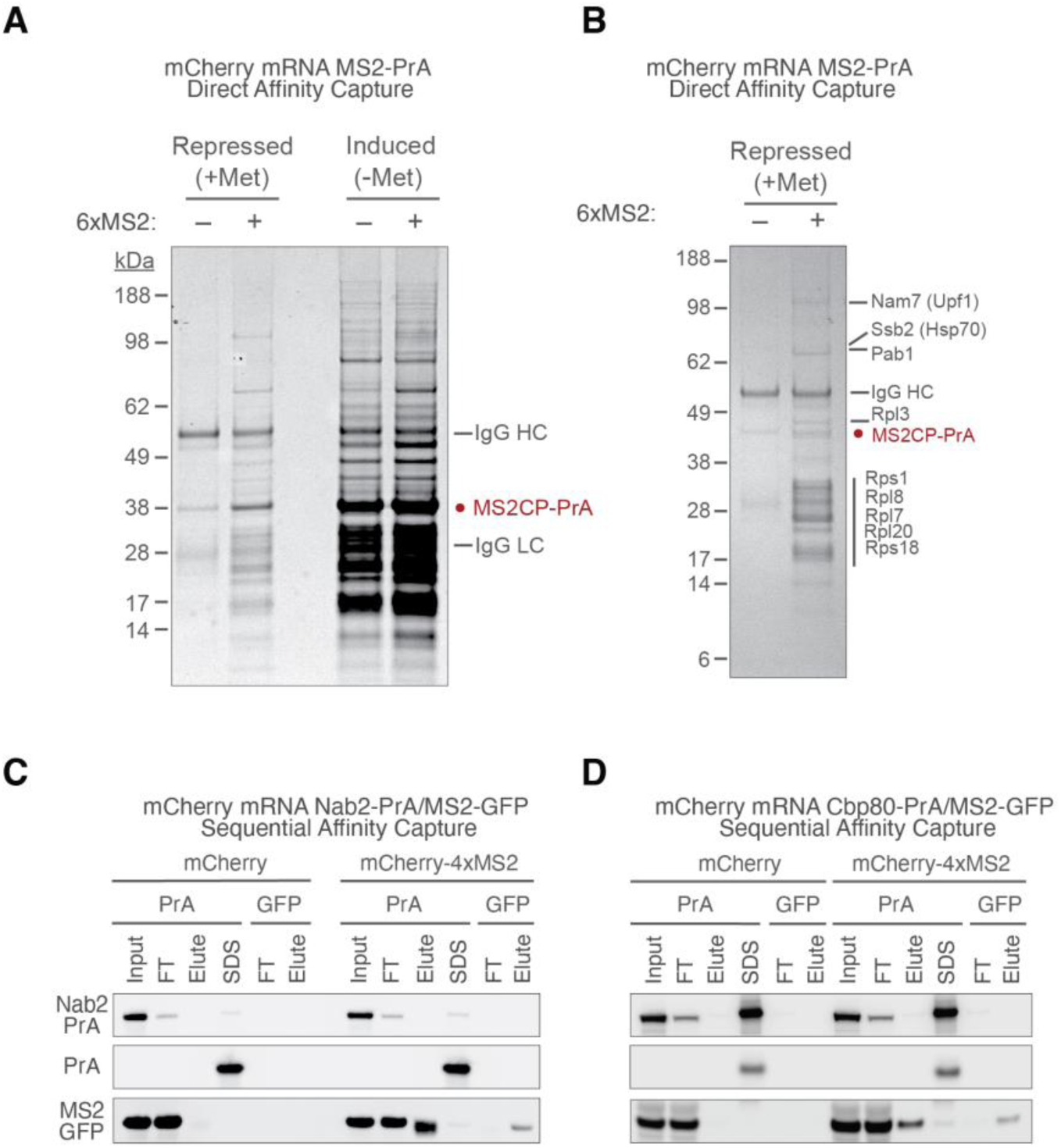
Validation and specificity of MS2-based mRNP isolations. **(A)** Direct, single-step isolations were performed of untagged or MS2-tagged mCherry RNA through capture of MS2CP-PrA expressed on a *MET3* promoter under repressive or permissive media methionine concentrations, indicating non-specific protein capture under high MS2-CP expression. **(B)** Indicated gel regions of a single-step *MET3*-repressed isolation of mCherry-6xMS2 through MS2CP-PrA were excised and analyzed by MALDI. Bands are labeled with corresponding protein IDs, indicating predominance of cytoplasmic RNA-associated factors. (C, D) Western blots of the PrA and GFP tags across the sequential isolation of untagged or 4xMS2-tagged mCherry, with PrA-tagged Nab2p (C) or Cbp80p (D), confirming step yields and strict 4xMS2-dependence of MS2CP-GFP recovery.

**Supplemental Figure 10.**
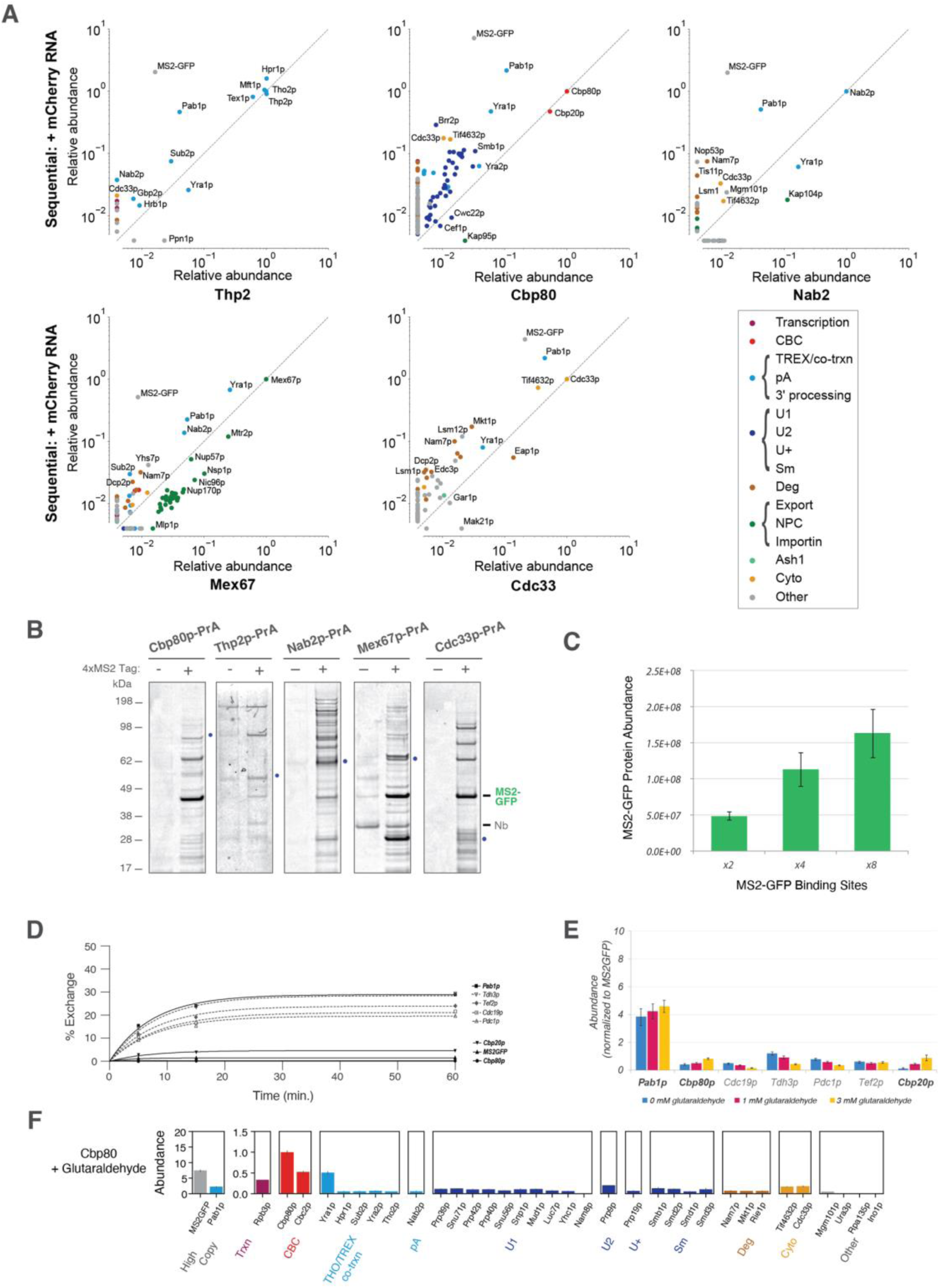
Characterization of sequential mRNP isolations across baits. **(A)** Comparison of glutaraldehyde-stabilized sequential isolations with single-step isolations of the same bait, showing MS2-step enrichment of transcript-associated proteins. **(B)** Coomassie-stained SDS-PAGE of elutions from sequential MS2CP-GFP/RNP-PrA isolations of untagged or 4xMS2-tagged mCherry transcripts across all bait strains. **(C)** Increase of MS2CP-GFP signal with MS2 hairpin number (2, 4, 8 hairpins) was determined by label-free MS quantification of elutions from sequential isolations of mCherry RNA with the indicated MS2 tag. **(D)** Post-lysis protein exchange measured by heavy/light mixing over time. Heavy isotopically labeled lysate was added to light (unlabeled) sequentially isolated MS2CP-GFP/Cbp80p complex on beads and incubated for the indicated time. Exchange of heavy/light protein was quantified for representative mRNA-associated proteins (black, bold) and common contaminant (gray). **(E)** Complex stabilization by the addition of the indicated concentrations of glutaraldehyde to cell grindate after resuspension, quenched after 5 min. incubation. Label-free MS quantification is shown for representative known interactors (black, bold) or contaminants (gray). **(F)** Relative protein copy number in Cbp80p-PrA/mCherry-4xMS2 mRNPs after 3 mM glutaraldehyde stabilization, normalized to Cbp80p; high-copy proteins on a separate scale and proteins below 5% of Cbp80p omitted—same layout as Figure 5A.

